# Comprehensive analysis of PNA-based antisense antibiotics targeting various essential genes in uropathogenic *Escherichia coli*

**DOI:** 10.1101/2022.02.21.481268

**Authors:** Linda Popella, Jakob Jung, Phuong Thao Do, Lars Barquist, Jörg Vogel

**Affiliations:** Institute of Molecular Infection Biology (IMIB), University of Würzburg, D-97080 Würzburg, Germany; Helmholtz Institute for RNA-based Infection Research (HIRI), Helmholtz Centre for Infection Research (HZI), D-97080 Würzburg, Germany; Faculty of Medicine, University of Würzburg, D-97080 Würzburg, Germany

## Abstract

Antisense peptide nucleic acids (PNAs) that target mRNAs of essential bacterial genes exhibit specific bactericidal effects in several microbial species, but our mechanistic understanding of PNA activity and their target gene spectrum is limited. Here, we present a systematic analysis of PNAs targeting eleven essential genes with varying expression levels in uropathogenic *Escherichia coli* (UPEC). We demonstrate that UPEC is susceptible to killing by peptide-conjugated PNAs, especially when targeting the widely-used essential gene *acpP*. Our evaluation yields three additional promising target mRNAs for effective growth inhibition, *i.e.*, *dnaB, ftsZ,* and *rpsH*. The analysis also shows that transcript abundance does not predict target vulnerability and that PNA-mediated growth inhibition is not universally associated with target mRNA depletion. Global transcriptomic analyses further reveal PNA sequence-dependent but also -independent responses, including the induction of envelope stress response pathways. Importantly, we show that the growth inhibitory capacity of 9mer PNAs is generally as effective as their 10mer counterparts. Overall, our systematic comparison of a range of PNAs targeting mRNAs of different essential genes in UPEC suggests important features for PNA design, reveals a general bacterial response to PNA conjugates and establishes the feasibility of using PNA antibacterials to combat UPEC.

## INTRODUCTION

Short antisense oligomers (ASOs) designed to target mRNAs of essential bacterial genes have emerged as attractive species-specific programmable RNA antibiotics and have been successfully applied against several bacterial species (1–5). Popular ASOs include peptide nucleic acids (PNAs), which harbor a pseudo-peptide backbone that links the four natural nucleobases. This chemical structure confers resistance towards nucleases and proteases. Since the neutral backbone also prevents charge repulsion, PNAs have strong binding affinities to complementary target sequences (4, 5). These properties allow the use of short oligomers – usually in the range of 9-12mers, to block a target (6–8). The capacity of PNA to base-pair with complementary sequences makes them *bona fide* antisense drugs, with the ability to inhibit gene expression both on the transcriptional (9–12) and posttranscriptional level (6,8,13,14). PNAs are generally designed to be complementary to the translation start site of a target mRNA and sterically block ribosome binding and initiation of translation (1,6,13). In order to facilitate intrabacterial delivery, PNAs are commonly conjugated to short, usually cationic or amphiphilic cell-penetrating peptides (CPPs) that facilitate crossing of the bacterial envelope (5,15–17). CPPs possess different penetration efficiencies depending on the targeted bacterial species (15–17).

Guidelines for the design of antisense PNAs take into consideration mostly features of the PNA molecule itself, such as its optimal length, its target region, infrequent off-targeting, and avoiding self-complementarity or long G-stretches (1,6–8,13,18). However, these guidelines are based on the analysis of a small number of PNA target genes, and generally do not include investigation of the characteristics of the PNA target genes themselves. The latter is also true for a recent side-by-side analysis of various PNAs designed with a novel semi-automated pipeline, which focused mainly on growth inhibition as a hallmark of PNA activity (19). This prompted us to systematically investigate PNA activity with respect to the choice of the PNA target gene, the characteristics of the applied PNA, and the physiological consequences of PNA treatment. We chose uropathogenic *Escherichia coli* (UPEC) as a model organism, due to its clinical relevance. UPEC is the primary cause of urinary tract infections, with around 150 million cases each year, and the emergence of antibiotic resistant UPEC strains creates the need for the development of alternative treatment options (20–24).

Firstly, we aimed to define the characteristics of an effective PNA target. One widely-used PNA target is the essential gene *acpP*, which is involved in fatty acid biosynthesis (25). PNA-mediated silencing of *acpP* mRNA, using concentrations similar to conventional antibiotics, causes strong bactericidal effects in several bacterial species, including *E. coli* and *Salmonella enterica* (6– 8,26,27). Although additional PNA targets that mediate bacterial growth inhibition have been identified, as reviewed elsewhere (4, 5), *acpP* is the most promising one due to its exceptional antibacterial activity. What makes *acpP* such an efficient target is still unclear; in fact, the high cellular abundance of the *acpP* mRNAs (28) makes it an unlikely target, given the unfavorable PNA/mRNA stoichiometry. However, the effect of target mRNA abundance on PNA activity has not been extensively investigated yet, and we therefore explored this hypothesis by designing PNAs targeting eleven essential genes with varying expression levels, including *acpP*.

Secondly, growth inhibition is the primary measure of efficacy of essential gene silencing by PNAs, generally measured using the endpoint-driven minimal inhibitory concentration (MIC) assay. However, growth inhibition does not provide information on wider PNA-triggered cellular responses. We therefore recently employed RNA-sequencing (RNA-seq) as a readout for how PNAs affect target mRNA level, and how PNA treatment globally reshapes the transcriptome in *Salmonella* (26). Although PNAs are designed to block translation of their target , there is growing evidence that PNAs also cause target mRNA depletion (18,26,27,29–31). However, PNA-triggered target mRNA decay and its relation to the efficacy of growth inhibition have not been systematically analyzed yet. Additionally, an analysis of the global transcriptomic landscape triggered by PNAs targeting different mRNAs to identify common responses is currently missing.

Finally, we focused on one important aspect which strongly influences the *in cellulo* antisense efficacy of a PNA, namely its length. Specifically, while short PNA molecules are taken up by the cell more easily, long PNA molecules are thought to have better binding affinities (6– 8,32–34). In *E. coli*, optimal PNA length has been proposed to be 10 nucleobases, but this is mostly based on the analyses of two genes, *acpP* (7) and *lacZ* (8). Currently, it has not been comprehensively tested whether this PNA length optimum is target gene-dependent or universally valid.

In the present study, our investigation of eleven essential genes identifies several promising new targets, while also reinforcing the unique antibacterial activity of *acpP* PNA. Further, we demonstrate that target gene abundance is not predictive of PNA target susceptibility. Our global RNA-seq analysis reveals that PNA-mediated target mRNA depletion is not a universal trait and is not essential for the antibacterial activity of a PNA. This analysis also identified common transcriptome-wide responses upon PNA treatment, such as induction of envelope stress response pathways. Finally, we show that 9mer PNAs, but not 8mers, are at least as efficient in growth inhibition as their commonly used 10mer counterparts. We consider this an important finding in the quest to reduce the off-target potential of antisense PNAs. In summary, our results provide insights into the druggable target gene spectrum, important aspects of PNA activity and global transcriptomic remodeling triggered by PNAs in UPEC.

## MATERIAL AND METHODS

### Bacterial strains, oligonucleotides and PNAs

The UPEC strain *E. coli* 536 was used throughout this study (internal strain number JVS-12054; NCBI GenBank: CP000247.1). The strain was streaked on Luria-Bertani plates and cultured in non-cation adjusted Mueller-Hinton Broth (MHB, BD Difco^TM^, Thermo Fisher Scientific), with aeration at 37°C and 220 rpm shaking.

Oligonucleotides were purchased from Eurofins (Eurofins Genomics) and dissolved in nuclease-free water. Sequences of all oligonucleotides used for amplifying the target region of the respective essential gene for generation of *green fluorescence protein* (*gfp)* fusion constructs (see “Overlap polymerase chain reaction (PCR) for generation of target gene-gfp fusion constructs”) are listed in Table S1.

Peptide-conjugated PNAs (PPNA), *i.e.* conjugated to (KFF)_3_K, (RXR)_4_XB, Tat or 2,3-diaminopropionic acid nonamer (Dap9), were obtained from Peps4LS GmbH. Sufficient quality and purity of these constructs was confirmed by mass spectrometry and HPLC (purity >98 %). PPNAs (Table S2) were dissolved in ultrapure water and heated at 55 °C for 5 min. Concentrations were determined by using a NanoDrop spectrophotometer (A _260 nm_; ThermoFisher) and calculated based on the extinction coefficient. PPNAs were stored at - 20 °C and incubated at 55 °C for 5 min before preparing the respective working dilutions. Low retention pipette tips and low binding Eppendorf tubes (Sarstedt) were used throughout.

### Minimal inhibitory concentration (MIC) assay

The broth microdilution method was applied for determination of MIC values according to the Clinical and Laboratory Standards Institute and a recently published protocol with a few modifications (35). Briefly, an overnight bacterial culture was diluted 1:100 in fresh MHB and grown to OD_600_ 0.5 (approximately 2-2.5 h). The obtained culture was diluted 1:2000 in fresh MHB to adjust a final cell concentration of approximately 10^5^ cfu/ml. Subsequently, 190 µl of the diluted bacterial solution was dispensed into a 96-well plate (Thermo Fisher Scientific). After adding 10 µl of 20x PPNA working solutions (ranging from 200 µM to 6.25 µM), or an equivalent volume of water as negative control, growth was monitored in a Synergy H1 plate reader (Biotek) by measuring the OD at 600 nm every 20 min with continuous double-orbital shaking (237 cpm) at 37°C for 24 h. The MIC was determined as the lowest concentration of PPNA, which inhibited visible growth in the wells (OD_600_ < 0.1).

### Killing time kinetics (bactericidal effects)

An overnight bacterial culture was diluted 1:100 in fresh MHB and grown to OD_600_ 0.5. The obtained culture was diluted 1:2000 in fresh MHB to adjust a final cell concentration of approximately 10^5^ cfu/ml. After pre-incubating the sample at 37 °C for 5 min, 10 µl of this diluted bacterial culture as well as serial dilutions (10^-1^, 10^-2^, 10^-3^) were directly spotted onto LB agar plates. Simultaneously, an adequate dilution was prepared and streaked on LB plates, serving as input condition for cfu determination. Further, 190 µl of the bacterial solution was transferred into 2 ml eppendorf tubes. Immediately, 10 µl of 20x PPNA solutions were added, adjusting the respective MIC. An equivalent volume of water was used as negative control, whereas a scrambled PPNA served as sequence-unrelated control. At the indicated time points post treatment, aliquots were used for cfu determination and spot assay.

### Overlap polymerase chain reaction (PCR) for generation of gfp fusion constructs

For the generation of target gene-*gfp* fusion constructs, oligonucleotides were designed to amplify the genomic region spanning nucleotides -40 to +51 relative to the translational start codon of each PNA’s target gene. This PCR step involved the attachment of the T7 promoter sequence to the 5’ end, to allow following transcription of the respective *gfp* fusion products, and the attachment of a 30 nucleotide *gfp*-overlap at the 3’ end, for PCR-aided fusion to *gfp*. A colony of UPEC 536 was resuspended in water and used as DNA template. In a separate PCR, *gfp* was amplified from pXG-10 plasmid (36) DNA and subsequently used for overlap PCR with each individual target gene region. All oligonucleotide sequences are listed in Table S1.

The PCR cycles for the primary amplification of target gene fragments and *gfp* were selected as follows: 98 °C for 1 min, 30 x 98 °C for 10 sec - 60 °C for 20 sec - 72°C for 20 sec, 72°C for 5 min, 10 °C ∞. Subsequently, the secondary overlap fusion PCRs were performed by using the oligonucleotides annealing at the 5’ end of the target gene and the 3’ end of *gfp*, with each 10-50 ng per PCR product as template. Conditions for this PCR program were set as follows: 98 °C for 1 min, 40 x 98 °C for 10 sec - 60 °C for 20 sec - 72°C for 40 sec, 72°C for 10 min, 10 °C ∞. Afterwards, overlap PCR products were purified with NucleoSpin Gel and PCR Clean-up (Macherey-Nagel) according to the manufacturer’s instructions and DNA concentration was quantified by using NanoDrop spectrophotometer (A _260 nm_).

### In vitro transcription

Templates for T7 RNA polymerase-driven transcription were produced via PCR as described above. After column-based purification, approximately 800 ng of DNA products were subjected to 20 µL *in vitro* transcription reactions according to the manufacturer’s instructions (MEGAscript T7 kit, Ambion/Thermo Scientific). After incubating the samples at 37 °C for 3 h and 45 min, 2 units (U) of Turbo DNase were added per reaction for additional 15 min. Subsequently, 115 µL water, 15 µL Ammonium Acetate Stop solution and 3 volumes of ethanol were added for RNA precipitation at - 80 °C overnight. After centrifugation, pellets were washed using 70 % ethanol and finally resuspended in ultrapure water. RNA concentration was measured with Qubit (Fisher Scientific). For verification of the expected product size and RNA integrity, RNA gels (6 % PAA, 7 M urea) were prepared with subsequent staining using StainsAll (Sigma-Aldrich).

### In vitro translation, titration of CPP-PNA-inherent inhibitory effects and western blotting

For *in vitro* translation of target gene-*gfp* fusion constructs, the PURExpress® In Vitro Protein Synthesis Kit (New England Biolabs, E6800L) was used as described in the instructions with minor modifications. To ensure a precisely adjusted final RNA concentration of 100 nM in all samples, *in vitro* transcribed RNA instead of T7 promoter-containing template DNA was used in all translation reactions. Briefly, the final volume of each *in vitro* translation reaction was set to 10 µL, each containing 4 µL Solution A, 3 µL Solution B, and 1 pmol of *in vitro* transcribed heat-denatured RNA to adjust a final concentration of 100 nM. After 2 h at 37 °C, the tubes were immediately placed on ice. For denaturation, protein samples were diluted in 1x reducing protein loading buffer (62.6 mM Tris–HCl pH 6.8, 2 % SDS, 0.1 mg/mL bromophenol blue, 15.4 mg/mL DTT, 10 % glycerol) and the tubes were incubated at 95 °C for 5 min.

To analyze whether sequence-specific or sequence-unrelated PPNAs are capable of inhibiting *in vitro* translation of target gene-GFP constructs, RNA samples were pre-incubated with defined PPNA solutions at 37 °C for 5 min. In particular, PPNA-inherent effects were titrated using the following conditions: 1 pmol of the respective RNA was *in vitro* translated in reactions containing 1 µM, 500 nM, 200 nM and 100 nM PPNA, corresponding to 10:1, 5:1, 2:1 and 1:1 ratios of PPNA:RNA. After this pre-annealing step, PURExpress® *in vitro* translation mix was added and samples were handled as described above.

For visualization of *in vitro* translated protein amounts, samples were separated on SDS-PAA (12 % PAA) gels, with subsequent semi-dry western blot transfer on polyvinylidene fluoride (PVDF) membranes. To verify equal loading, membranes were first stained with Ponceau S (Sigma-Aldrich) and then probed with 5 % skim milk (in 1x TBS-T) containing anti-GFP antibody (1:1,000; Sigma-Aldrich) overnight. After incubation with an HRP-conjugated secondary antibody (1:10,000; ThermoScientific) in 1x TBS-T, the membrane was incubated with a self-made developing solution and protein levels were detected using an ImageQuant LAS 500 (GE Healthcare Life Sciences). Images were processed and band intensities quantified using ImageJ (37).

### PPNA treatment of UPEC 536 and isolation of total RNA

Bacterial overnight cultures were diluted 1:100 in fresh MHB and grown to an OD_600_ of 0.5. Subsequently, obtained cultures were again diluted 1:100 in fresh MHB to adjust a cell concentration of approximately 10^6^ cfu/mL. After transferring 1.9 mL of the bacterial solution into 5 mL low-binding tubes (LABsolute), 100 µL of 20x PPNA stock solutions were added to reach a final concentration of 5 µM for all KFF-conjugated PNAs. In parallel, an equal amount of cells was treated with the respective volume of sterile nuclease-free water, which was used as solvent for the test compounds, and served as negative control. After incubating the samples for 15 min at 37 °C, RNAprotect Bacteria (Qiagen) was added according to the manufacturer’s instructions. Following a 10-min incubation, cells were pelleted at 4 °C and 21,100 xg for 20 min. The supernatant was discarded and pellets were either directly used or stored at – 20 °C (< 1 day) for subsequent bacterial RNA isolation.

Total RNA was purified from bacterial pellets using the miRNeasy Mini kit (Qiagen) according to the protocol #3 described in Popella *et al.* (26). Briefly, cells were resuspended in 0.5 mg/mL lysozyme (Roth) in TE buffer (pH 8.0) and incubated for 5 min. Afterwards, RLT buffer supplemented with β-mercaptoethanol, and ethanol were added according to the manufacturer’s instructions. After sample loading, column wash-steps were performed according to the manual. RNA concentration was measured with a NanoDrop spectrophotometer.

### RNA-seq

For transcriptomic analyses, RNA samples were processed and subjected to RNA-seq at Core Unit SysMed (Julius-Maximilian-University Würzburg, Germany). Briefly, samples were DNase-treated and quality was checked on a bioanalyzer (RNA chip Agilent). Then, RNA was subjected to cDNA library preparation using the Corall kit (Lexogen) and including the depletion of ribosomal RNA (RiboCop-META kit, Lexogen) according to the manufacturer’s instructions. Afterwards, library samples were pooled in equimolar amounts and quality was verified using a bioanalyzer (DNA chip Agilent). The cDNA pools were sequenced using the NextSeq 500 system (HighOutput flow cell, 400 M, 1x 75 cycle single-end; Illumina).

### Quantification of RNA-seq data

RNA-Seq data analysis was performed similarly to our previous analysis of PNA activity in *S. enterica* serovar Typhimurium (26). Small RNAs (sRNAs) were added to the UPEC annotation file. Specifically, sRNA sequences of *E. coli* str. K-12 substr. MG1655 were aligned to the UPEC genome using BLASTN (v2.9.0), and the alignment match with the lowest E-value was added as an sRNA to the UPEC annotation. All added sRNAs had more than 89% nucleotide identity and the largest number of sequence mismatches was 4. Raw RNA-Seq reads were trimmed, filtered and mapped to the UPEC genome (*E. coli* 536, CP000247.1) (38). BBduk was used to remove adapter sequences and trim nucleobases with Phred quality scores below 10. Reads were then mapped against the UPEC genome using BBMap (v38.84) and the resulting alignments were quantified against both coding sequences and sRNAs using the featureCounts methods of the Subread (2.0.1) package (39).

### Normalization and differential expression analysis

R/Bioconductor packages were used for the downstream RNA-Seq analysis. Raw read counts from all conditions were used with edgeR (v3.34.1) for differential expression analysis (40). Prior to analysis, reads which did not meet the threshold of 1.48 counts per million (CPM) in at least 10 libraries were filtered out. The cutoff was set as 10/L, where L is the minimum library size across all samples in millions, as proposed in (41).

The remaining libraries were normalized by the trimmed mean of M values (TMM) normalization (42). Then, libraries were assessed for batch effects using principal component analysis (PCA) plots. A consistent batch effect was observed between different sequencing runs. As described in the edgeR manual (43), batch effects were corrected by adding a batch variable to the design matrix before differential expression analysis. Quasi-likelihood dispersions were estimated using the glmQLFit function. Then contrasts were created and used to test for differential expression with the glmQLFTest function.

Features with absolute fold changes >2 and false discovery rate (FDR) adjusted *P*-values (44) < 0.001 were considered differentially expressed. The results were plotted as heatmaps using the ComplexHeatmap (v2.8.0) package.

### KEGG pathway analysis

For assigning genes to KEGG pathways (45), the R package KEGGREST (1.32.0) was used. Additionally, gene sets of regulons were added. To assign genes to regulons, the *E. coli* K12 RegulonDB (v10.9) was parsed and genes appearing in both UPEC and K12 were assigned to regulons (46). Rotation gene set testing (FRY version of ROAST gene set testing, (47)) was applied to assess enrichments of pathways and regulons in differentially expressed genes. Gene sets with >10 genes and an FDR-adjusted *P*-value <0.001 (marked with an asterisk, *) were visualized in Figure 7, the color indicating the median log_2_ FC of the respective gene set. Figure 7 shows all gene sets which rank among the top 10 significant gene sets with the lowest *p-*value for at least one sample.

### Prediction of PNA/RNA melting temperature

Since there is no reliable bioinformatic algorithm for prediction of the melting temperature (Tm) of PNA/RNA duplexes currently available, we used the MELTING package to predict Tms for the corresponding RNA/RNA duplexes (48). We used our predicted values as estimation of the actual PNA/RNA Tm, which are known to be higher compared to the respective RNA/RNA duplexes (49). As proof of concept, we verified our Tm prediction based on previously published experimental Tms of PNA/RNA duplexes (7).

## RESULTS

### acpP PNA conjugated to (KFF)_3_K, (RXR)_4_XB, or Tat peptides inhibits growth of UPEC 536

To explore the application of PNAs to combat UPEC (Figure 1A), we took advantage of a commonly used *acpP* PNA with proven efficacy in numerous bacterial species (7,18,26,27,29,30,50,51). The PNA has full complementarity to 10 nucleotides located around the start codon of *acpP*, a site that is highly conserved in several γ-proteobacteria including the pyelonephritis isolate UPEC 536, widely used as model for extraintestinal *E. coli* (38) (Figure 1B-C).

**Figure 1.**
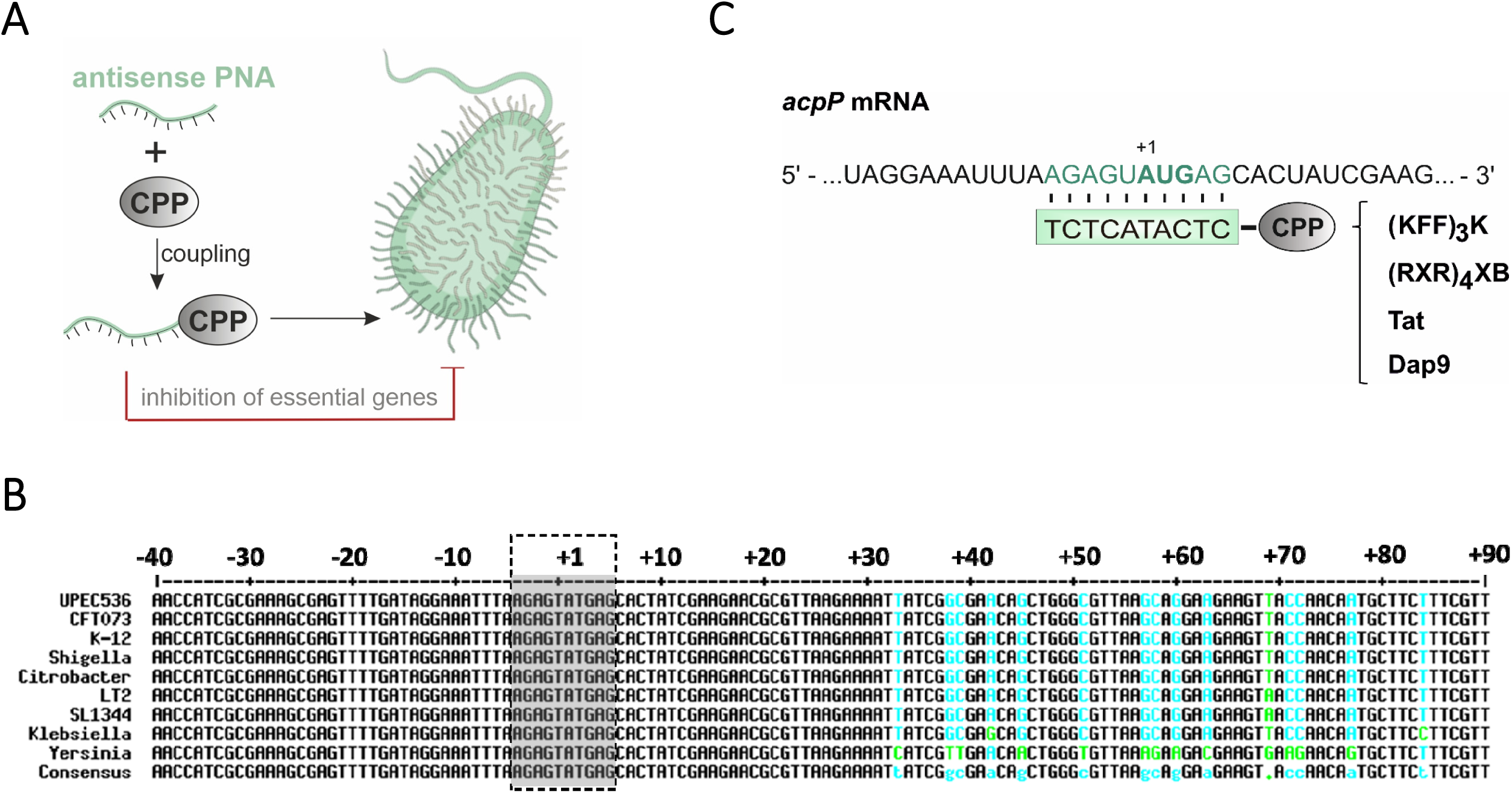
Schematic illustration of the applied PNA technology and the PNA target region of the conserved essential gene, *acpP*. **(A)** An antisense PNA is coupled to a CPP to facilitate intrabacterial delivery. When targeting essential bacterial genes, CPP-PNA conjugates can serve as potent antimicrobial molecules. Parts of this image have been created with Smart (Servier medical art). **(B)** Multiple sequence alignment (created with http://multalin.toulouse.inra.fr/multalin/; (116)) of the essential gene *acpP* including the following *γ-*proteobacteria: UPEC 536 (CP000247.1), *E. coli* CFT073 (CP051263.1), *E. coli* K-12 (CP032667.1), *Shigella dysenteriae* (CP000034.1), *Citrobacter rodentium* (CP038008.1), *Salmonella enterica* subsp. enterica serovar Typhimurium str. LT2 (AE006468.2), *Salmonella enterica* subsp. enterica serovar Typhimurium str. SL1344 (FQ312003.1), *Klebsiella pneumoniae* (CP003200.1), and *Yersinia pestis* (NC_003143.1). A defined section (-40 to +90 nt) including the region around the PNA binding site (grey box) is shown. The following color-code has been applied: perfect consensus black, varying alignment columns cyan (nucleotide substitution green). **(C)** Relevant region of *acpP* mRNA in UPEC 536, with the PNA target sequence shaded in green. The start codon (AUG) is shown in bold type. Below, the PNA sequence is shown (green box) with the different conjugated CPPs for delivery into UPEC ((KFF)_3_K, (RXR)_4_XB, Tat, Dap9).

Using UPEC 536, we determined the MIC of *acpP* PNA coupled to the CPPs (KFF)_3_K, (RXR)_4_XB, Tat, or 2,3-diaminopropionic acid nonamer (Dap9), which have previously been shown to mediate efficient bacterial permeation (16, 26) (Figure 2). We included scrambled PNA and peptide only controls in our analyses to differentiate between sequence-independent effects triggered by either the PNA module or by the carrier peptide itself. (KFF)_3_K-, (RXR)_4_XB-and Tat-coupled *acpP* PNAs all lead to efficient *in vitro* growth inhibition, with (KFF)_3_K and (RXR)_4_XB conjugated *acpP* PNA being more potent (MIC of 1.25 µM) than Tat-coupled *acpP* PNA (MIC 5 µM) (Figure 2, left panels). However, while (KFF)_3_K- or Tat-conjugated scrambled PNAs or the respective ‘peptide onl’ controls did not cause substantial growth inhibition, the (RXR)_4_XB compounds all impaired UPEC 536 growth. Thus, the (RXR)_4_XB peptide itself seems to possess antibacterial activity against UPEC 536 (MIC of 10 µM). On the other hand, the Dap9-conjugated *acpP* 10mer PNA did not mediate substantial growth inhibition of UPEC 536 (MIC > 10 µM; Figure 2D). Based on these data, we conclude that the (KFF)_3_K peptide is a potent carrier for PNA delivery into UPEC 536 and we therefore used it in all subsequent analyses.

**Figure 2:**
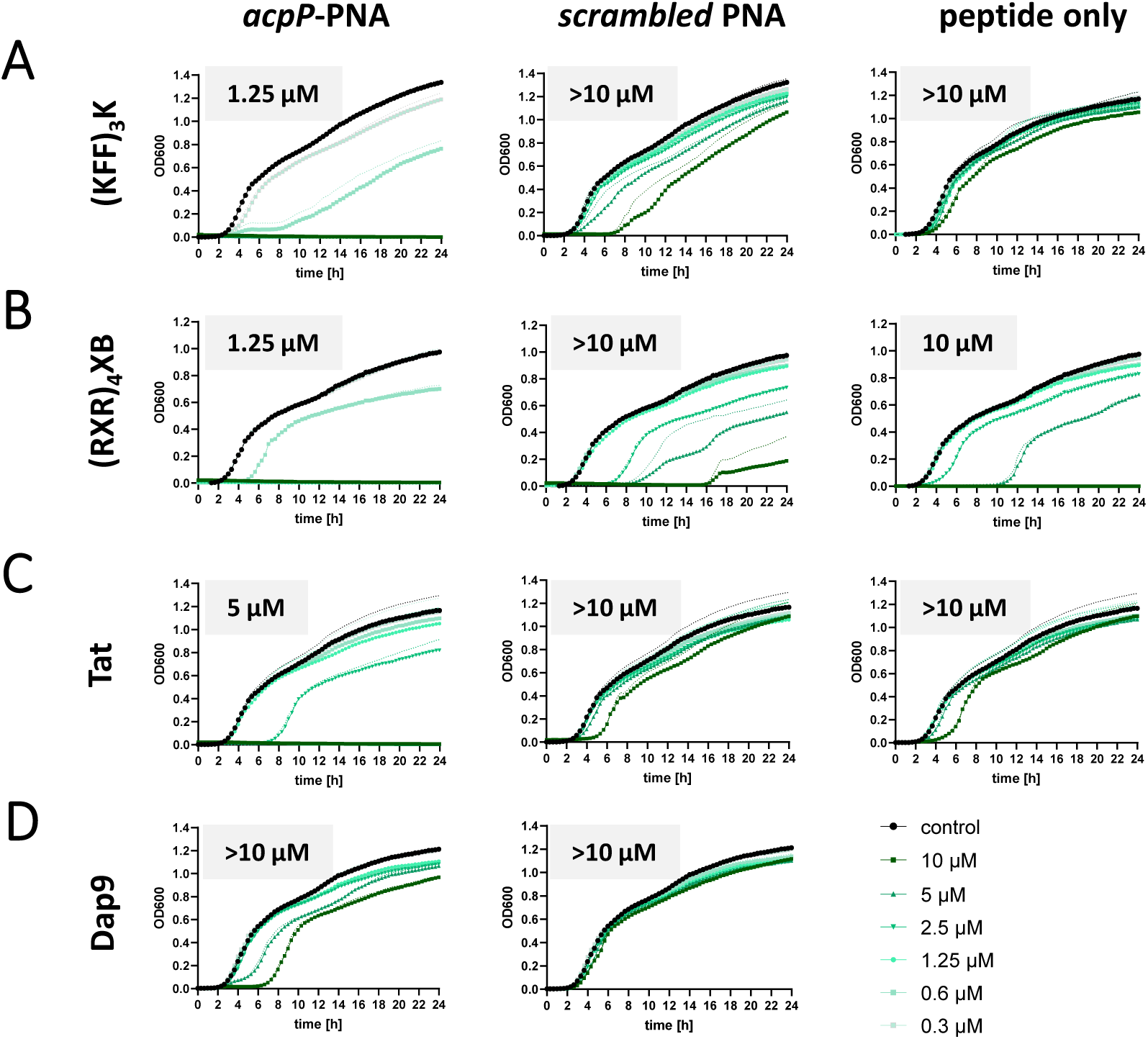
An antisense PPNA targeting *acpP* shows potent growth inhibitory capacity against UPEC. Determination of growth kinetics and MICs of UPEC 536 upon exposure to *acpP*-specific (left panels) and scrambled control (middle panels) PNAs, conjugated to **(A)** (KFF)_3_K, **(B)** (RXR)_4_XB, **(C)** Tat or **(D)** Dap9 peptide. **(A-D)** UPEC 536 (10^5^ cfu/ml) were exposed to gradually decreasing concentrations (starting from 10 µM) of each PPNA or peptide. Treatment with an equal volume of water served as control (black curves). **(A-C)** CPPs (right panels) were used as PNA-independent controls. Growth is depicted as OD_600_ (y-axis) and was monitored over 24 h (x-axis). Each experiment was performed two to four times. Curves represent the mean of all biological replicates and dashed lines indicate standard error of the mean.

### (KFF)_3_K-acpP mediates bactericidal effects in UPEC 536

To clarify if the observed growth inhibitory capacity of (KFF)_3_K-*acpP* was an indication of bacteriostatic or bactericidal effects, UPEC 536 cells were challenged with the MIC of 1.25 µM for different time intervals, with water or (KFF)_3_K-*acpP*-scrambled serving as controls (Figure 3). For all conditions, spot assays with serial dilutions (Figure 3A) as well as colony forming units (cfu) per mL (Figure 3B) were used to measure bacterial survival. As expected, scrambled (KFF)_3_K-*acpP* did not affect the abundance of living bacterial cells compared to the water control within 24 h. In sharp contrast, treatment of UPEC 536 with (KFF)_3_K-*acpP* strongly decreased the number of cfu, evident from 30 minutes post treatment (mpt) onwards. A plateau of the bactericidal effect was observed from 60 to 120 mpt, revealing a 1 log reduction in living cells. This effect was even more pronounced after 24 h of treatment, showing a 2-3 log decrease in cfu/mL. These data demonstrate that (KFF)_3_K-*acpP* acts as a potent antibacterial agent that possesses sequence-specific activity against UPEC 536.

**Figure 3:**
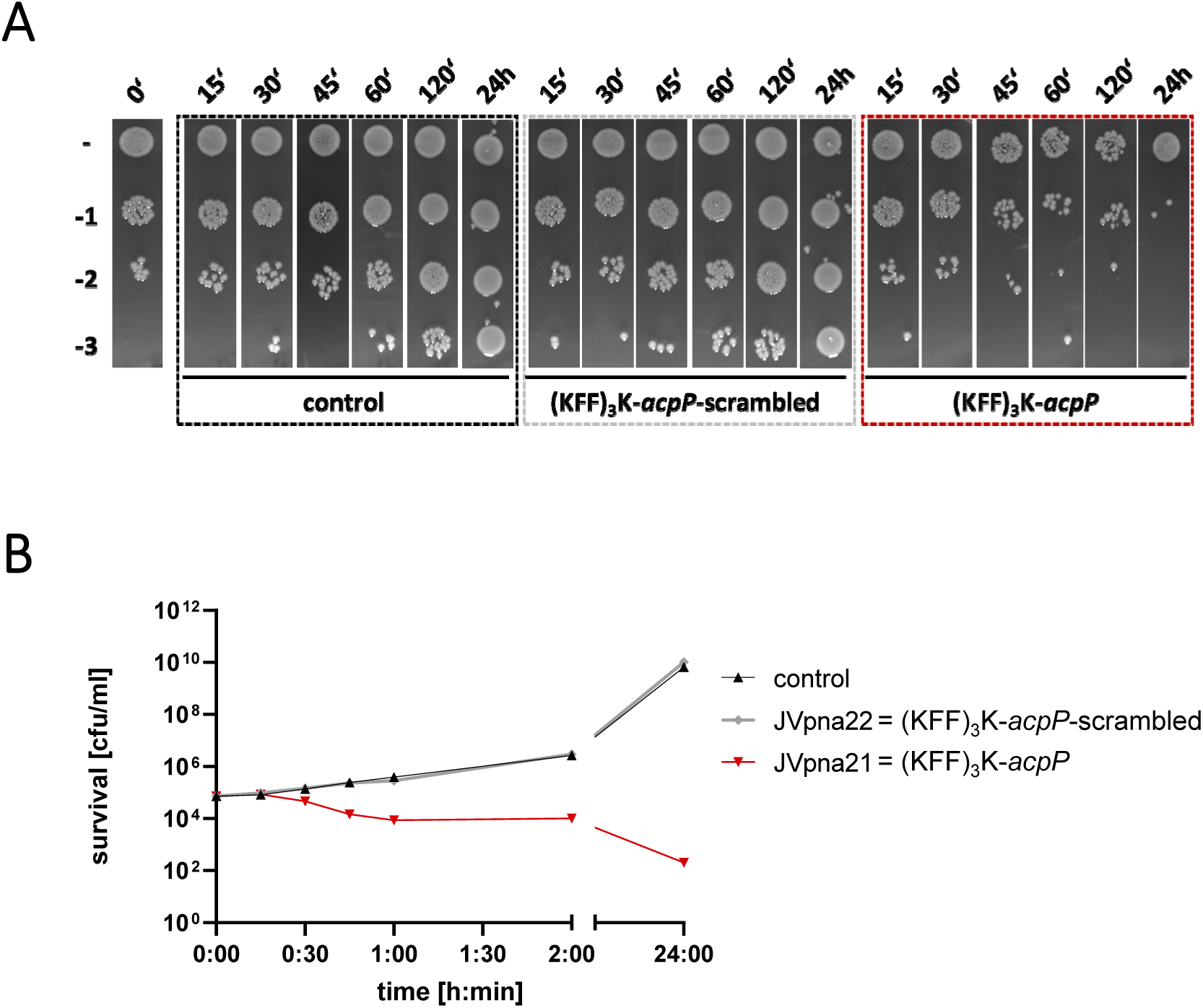
An antisense PNA targeting *acpP* shows antibacterial activity against UPEC 536 at 1x MIC. Bactericidal effects of (KFF)_3_K-*acpP* (red) were determined at 1x MIC (1.25 µM) against UPEC 536 (10^5^ cfu/ml) during a 24 h time course. (KFF)_3_K*-acpP*-scrambled (grey) was used as control PNA, whereas exposure to an equal volume of water served as additional control (black). After the indicated time points, aliquots of each condition were subjected to **(A)** spot assays or **(B)** cfu determination on LB plates to investigate the amount of viable cells. **(A)** Serial dilutions (10^-1^ to 10^-3^) were prepared, as indicated on the left. **(B)** Curves represent the mean of the biological replicates. The experiments were performed twice.

### (KFF)_3_K-acpP inhibits in vitro synthesis of AcpP^1-17^-GFP

Next, we investigated whether (KFF)_3_K-*acpP* inhibits *acpP* translation, as would be expected from the sequence-complementarity of the PNA to the translational start site of its target mRNA. To this aim, we applied a cell-free *in vitro* translation system. Specifically, the genomic region spanning - 40 to +51 of the *acpP* mRNA relative to the start site, including the PNA target region, was fused to *gfp* (*acpP*^91^*::gfp*). After PCR amplification and *in vitro* transcription, the *acpP^91^::gfp* construct was translated *in vitro* and AcpP^1-17^-GFP protein abundance was determined via western blotting (Figure 4). Titration of the (KFF)_3_K-*acpP* construct from 10:1, 5:1, 2:1, and 1:1 molar ratios to *acpP^91^::gfp* RNA showed that a 5-fold molar excess of (KFF)_3_K-*acpP* over *acpP^91^::gfp* RNA was already sufficient to trigger an 82 % reduction in translation efficiency on average, compared to the water-treated control (Figure 4A). We also included (KFF)_3_K-*acpP*-scrambled control at 10-molar excess over *acpP^91^::gfp* RNA, to confirm sequence-specific inhibition of translation *in vitro*. While (KFF)_3_K-*acpP* substantially inhibited *in vitro* synthesis of AcpP^1-17^-GFP, the scrambled peptide-conjugated PNA (PPNA) construct did not affect translation efficacy (Figure 4B). These results demonstrate that (KFF)_3_K-*acpP* is capable of inhibiting cell-free translation of a truncated *acpP*-target fused to GFP.

**Figure 4:**
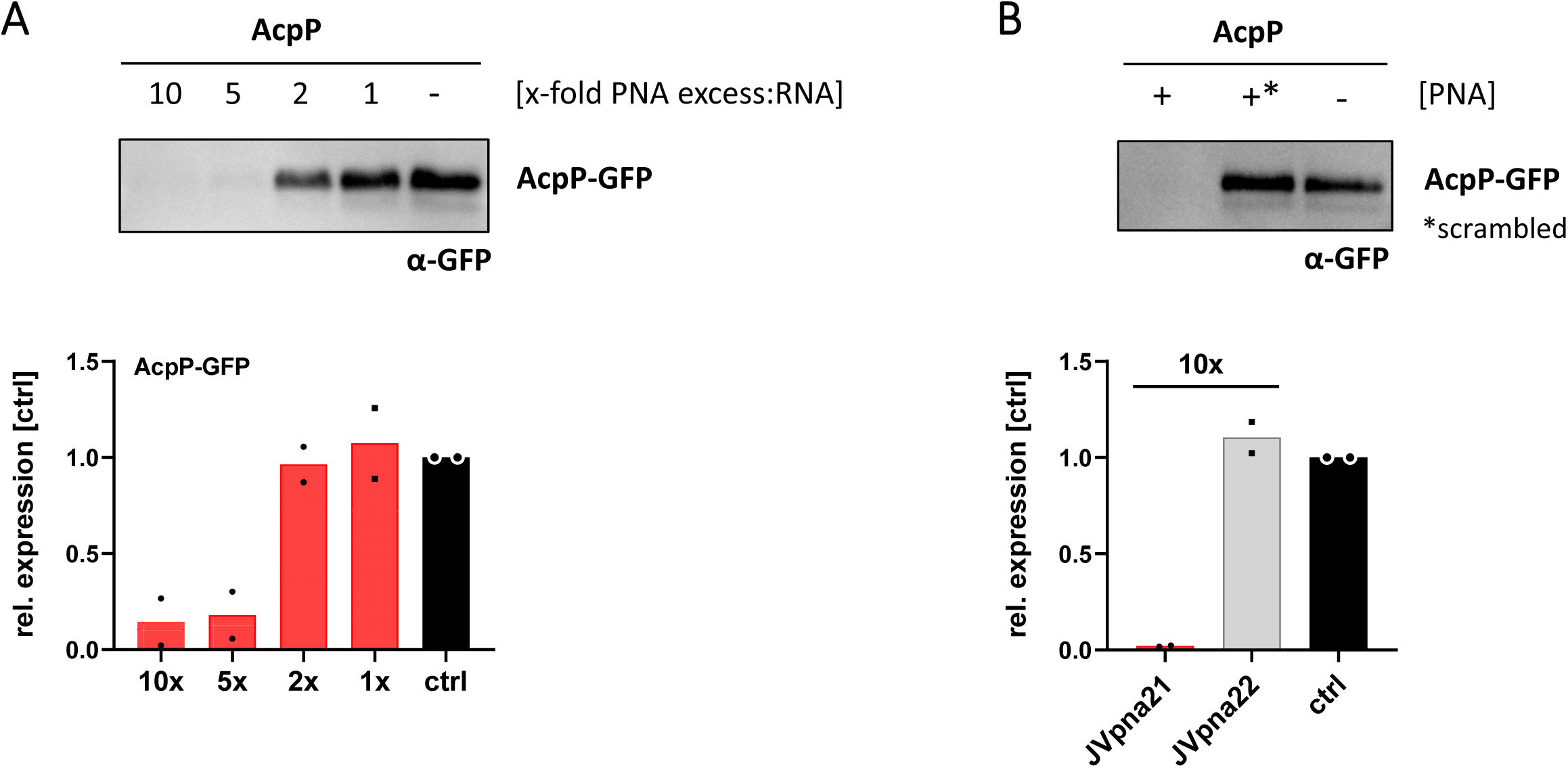
An antisense PNA targeting *acpP* inhibits translation of its target *in vitro*. *In vitro* transcribed *acpP::gfp* RNA was subjected to *in vitro* translation assays. **(A)** Titration of the molar excess of (KFF)_3_K-*acpP* over template mRNA was performed to determine the inhibitory efficacy. As control, an equal volume of water was added (-, ctrl). Western Blotting was performed for detection of the AcpP^1-17^-GFP fusion (approximately 25 kDa) by using an anti-GFP antibody. Signal intensities were quantified using ImageJ and protein expression is shown relative to the “untreated” control. Black dots indicate individual protein expression levels of the experimental duplicates and bars show the mean. **(B)** *In vitro* translation was performed in the presence or absence of (KFF)_3_K-*acpP* (+; Jvpna21) or its scrambled control (+*; Jvpna22). A 10-molar excess of both PNA constructs over template RNA was applied. The experiments were performed two times.

### Expanding the essential gene spectrum for PPNA targeting of UPEC 536

In order to investigate if PNA-mediated antibacterial activity can be harnessed more widely and to clarify whether transcript abundance influences the efficacy by which PNAs deplete their target, we selected additional essential genes with differing expression levels for PNA-based targeting. We first ranked genes previously shown to be essential in a variety of *E. coli* strains (52–54) by expression level, and chose ten target genes covering the range of observed expression (Figure 5A). We designed 10mer antisense PNA sequences to target the translational start sites of these genes in a window spanning nucleotides -6 to +7. With the exception of the PNAs for *pyrH* or *rplS*, the PNAs we designed do not have predicted off-target sites within the translation initiation regions of other genes. We then performed *in vitro* translation assays in the presence or absence of the respective sequence-specific PPNA for each selected target gene as described above for *acpP* (Figure 4). *In vitro* translation of *dnaB* and *nusG* was unsuccessful in our assay conditions, and we therefore excluded both genes from this analysis. With the exception of *rplS*, all PPNAs strongly decreased the protein abundance of the respective target fusion (Figure 5B). Since the PPNA/RNA duplex stability affects the antisense efficacy of a PNA (7,8,32–34), we analyzed whether the stability also affects the inhibition of translation *in vitro*. Our analysis reveals a direct association of the predicted melting temperature*, i.e.*, PPNA/RNA duplex stability, with the capacity of a PPNA to inhibit translation of its target *in vitro* (Figure 5C).

**Figure 5:**
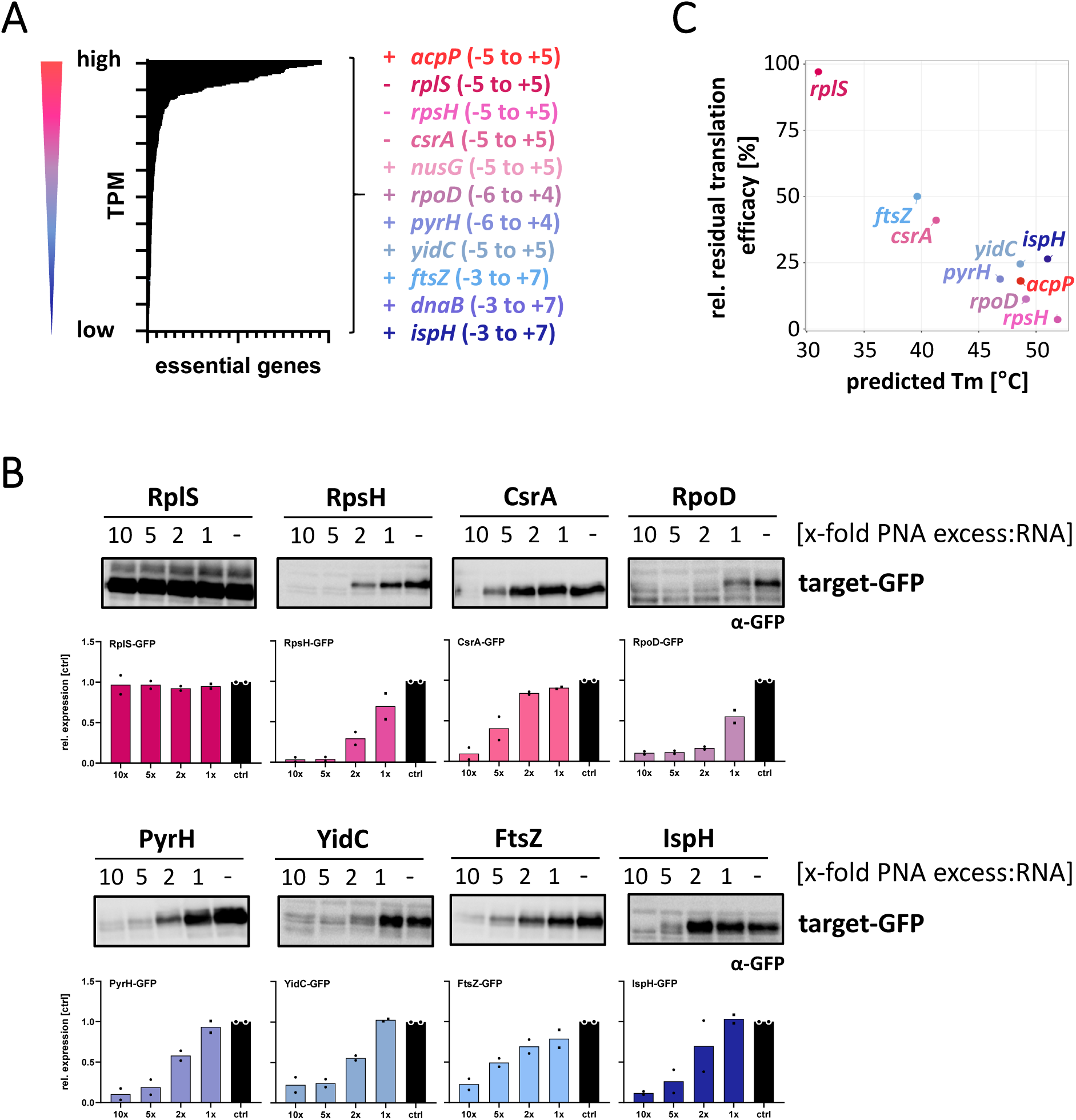
Expanding the target spectrum of essential gene-specific antisense PNAs in UPEC. **(A)** All essential genes, annotated in *E. coli* (52–54), were ranked according to their expression levels in our UPEC 536 RNA-seq data set (untreated control samples, averaged from three independent biological replicates). High to low expression levels are indicated by red to blue coloring. Essential gene candidates for PNA targeting were selected to cover a broad range of mRNA abundances (indicated on the right side). The coding strand of each gene is indicated with (+) or (-). The target window is indicated as nucleotide positions relative to the start codon. **(B)** *In vitro* transcribed *gfp-*fusion RNAs for each essential gene were subjected to *in vitro* translation assays, in the presence or absence of the cognate PPNA. The capacity of the tested PPNAs to inhibit *in vitro* translation was analyzed in 10:1, 5:1, 2:1 to 1:1 molar ratios of PPNA:RNA. An equal volume of water was added to serve as control condition (-, ctrl). Samples were subjected to Western blotting and membranes were probed with an anti-GFP antibody to determine the expression levels of the GFP fusion proteins (approximately 25 kDa). Signal intensities were quantified using ImageJ. Protein expression levels are shown relative to the respective control sample. Black dots indicate individual protein expression levels of the experimental duplicates and bars show the mean. Each experiment was performed twice. **(C)** Averaged residual translation efficacy, relative to the respective control condition, at 5-molar excess of PNA over RNA, plotted against the predicted melting temperature (Tm, °C) of each PNA/RNA duplex.

### Target mRNA abundance does not dictate the growth inhibitory capacity of a PPNA

Next, we determined the MIC of each PPNA and observed that 7 out of 10 tested (KFF)_3_K-conjugated 10mer PNAs are capable of inhibiting visible growth of UPEC 536 at concentrations ≤ 10 µM (Table 1). PPNAs targeting *yidC, csrA*, and *nusG* did not trigger complete growth inhibition, although the two former did cause growth reduction at 10 µM (Figure S1). These effects were not correlated with target gene abundance (Figure 6A).

**Figure 6:**
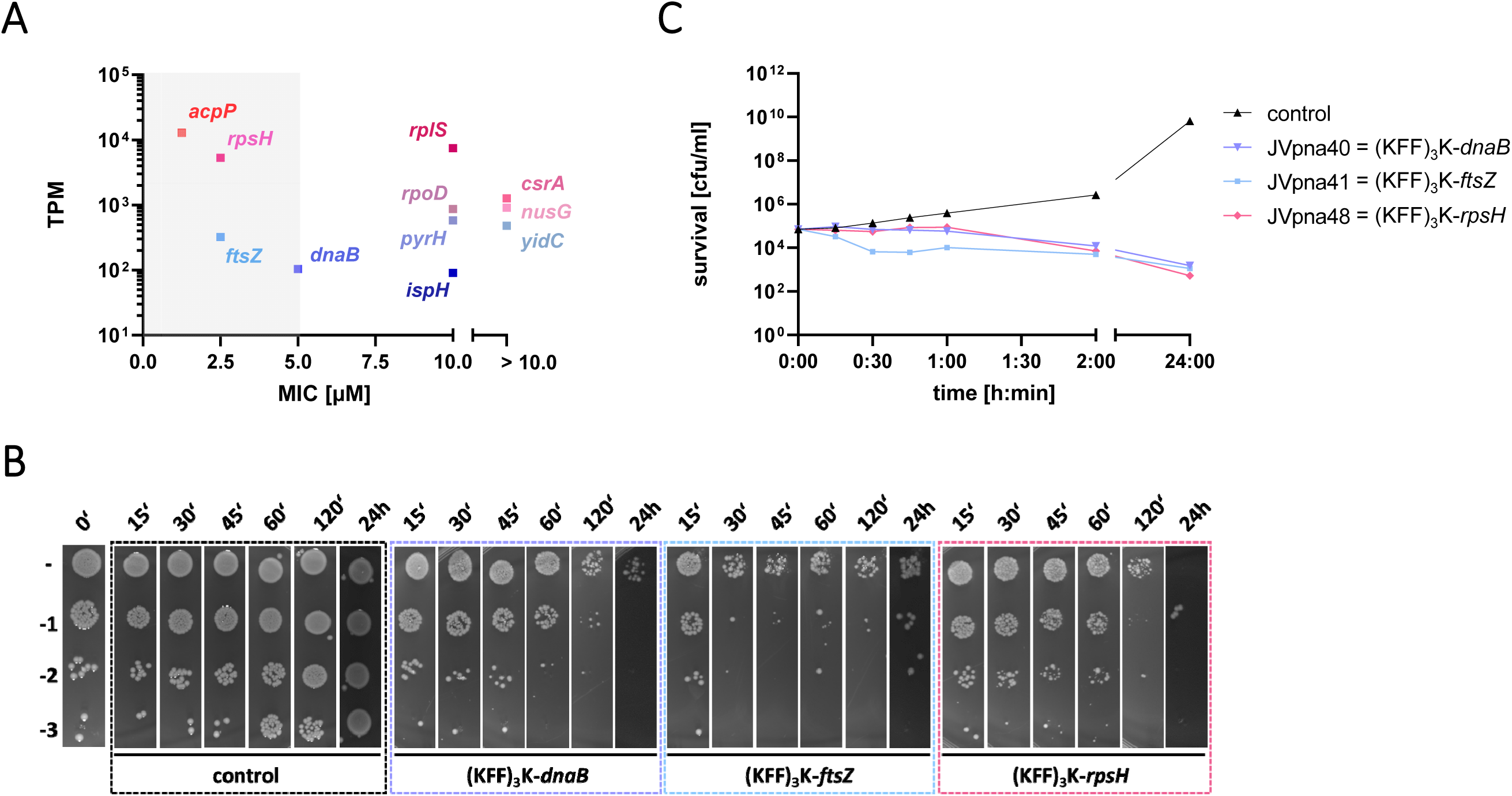
Antisense PPNAs targeting *dnaB, ftsZ,* and *rpsH* confer antibacterial activity against UPEC 536 at 1x MIC. **(A)** MICs for each 10mer PPNA, listed in Table 1, were plotted against the abundance (transcripts per million, TPM) of the respective PNA’s target. Grey shaded area highlights the four PPNAs with the strongest antibacterial efficacy against UPEC 536. **(B-C)** Bactericidal effects were determined for (KFF)_3_K-*dnaB* (5 µM), (KFF)_3_K-*ftsZ* (2.5 µM), and (KFF)_3_K-*rpsH* (2.5 µM) at 1x MIC against UPEC 536 (10^5^ cfu/ml). Exposure to an equal volume of water served as control (black). Samples were taken at the indicated time points post treatment. **(B)** Spot assays with serial dilutions (10^-1^ to 10^-3^) and **(C)** cfu evaluation on LB plates was performed to determine the number of viable cells. Curves represent the mean of the biological replicates. Each experiment was performed twice.

**Table 1:**
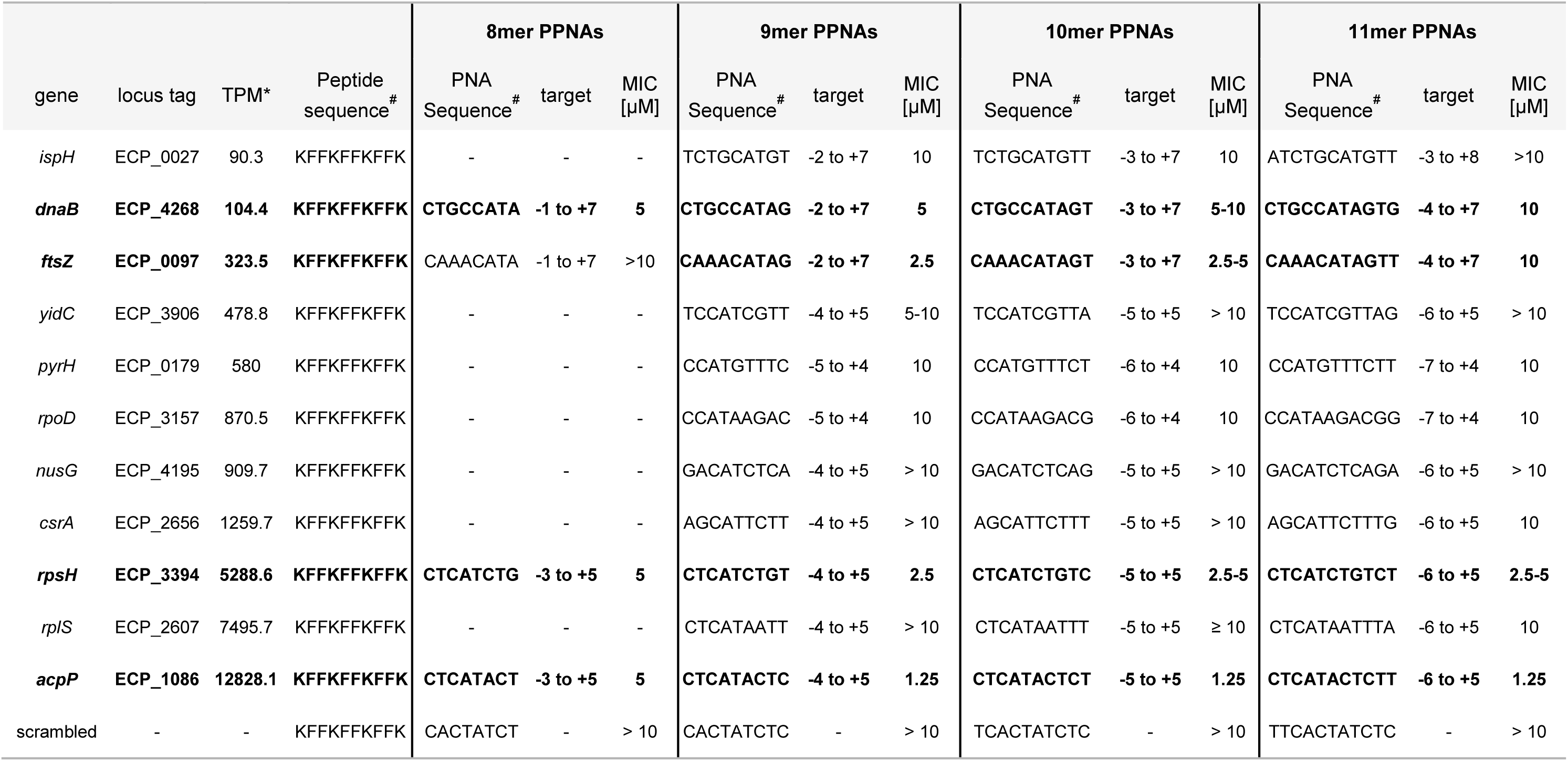
Analyzing the growth inhibitory capacity of 8-11mer PPNAs targeting selected low to high abundant essential genes. UPEC 536 cells (10^5^ cfu/ml) were exposed to serial dilutions of (KFF)_3_K-coupled 8mer, 9mer, 10mer or 11mer PPNAs and growth was monitored after 24 h. Minimum inhibitory concentrations (MICs) were determined. The PPNAs are ordered according to the target transcript abundance from low to high expression levels (transcript per million, TPM). * Averaged value from RNA-seq analyses of three untreated control samples. ^#^ Sequences are shown from N- to C-termini orientation.

For further characterization, we focused on PNAs targeting *ftsZ, dnaB* and *rpsH*, which showed MICs of less than 10 µM (Figure 6A). Notably, while PNAs targeting *ftsZ* have previously been shown to inhibit growth of *E. coli*, *Enterococcus faecalis* and *Acinetobacter baumannii* (29,50,55), *dnaB* and *rpsH* are novel PNA targets. We assayed bactericidal effects of these PPNAs against UPEC 536 at their MIC, via spot assays (Figure 6B) and cfu determination (Figure 6C). Similar to our observations for (KFF)_3_K-*acpP* (Figure 3), treatment with the three PPNAs targeting *dnaB, ftsZ* or *rpsH* caused reduction in bacterial survival, with (KFF)_3_K-*ftsZ* exhibiting an accelerated effect compared to both other constructs. All three tested PPNAs led to a 1 to 2 log reduction in cfu/mL evident from 60-120 min onward.

Overall, these results exclude target mRNA abundance as a key determinant for the growth inhibitory capacity of a PPNA in UPEC, consistent with a recent study in multidrug resistant *Enterobacteriaceae* (19). Surprisingly, the efficacy of *in vitro* inhibition of translation (Figures 4 and 5B) also does not correlate with PPNA activity against UPEC.

### Global transcriptomic analysis reveals specific and common (KFF)_3_K-PNA-mediated transcriptional changes

In order to investigate the global transcriptomic response of UPEC when challenged with different PPNAs and to test if the previously observed target transcript depletion (26) is a universal effect of PPNA activity, we performed RNA-seq analysis. Specifically, UPEC 536 was exposed to equimolar concentrations of the 10mer (KFF)_3_K-coupled PNAs listed in Table 1. After 15 min, cells were harvested and RNA was isolated for subsequent RNA-seq analyses. As a quality control, we evaluated the RNA-seq dataset by principal component analysis (PCA) and hierarchical cluster analysis (HCA) (Figure S2). After removing batch effects within our triplicates, we found that all biological replicates of one condition clustered together.

We initially performed a gene set enrichment analysis to identify globally regulated pathways (Figure 7). We included annotated KEGG pathways (45) and manually added the virulence and stress response-associated regulons GadX, PhoPQ, CpxR, SlyA, SoxS, MarA, Rob and Fis (marked in blue) retrieved from RegulonDB (46). Across the analyzed PPNAs, we observed a significant induction (FDR-adjusted *P*-value < 0.001) of the cationic antimicrobial peptide resistance (CAMP), GadX (note: for (KFF)_3_K-*ftsZ* below the significance threshold), PhoPQ and CpxR pathways in PPNA-treated UPEC 536. This indicates that PPNAs trigger several common stress response pathways in UPEC.

**Figure 7:**
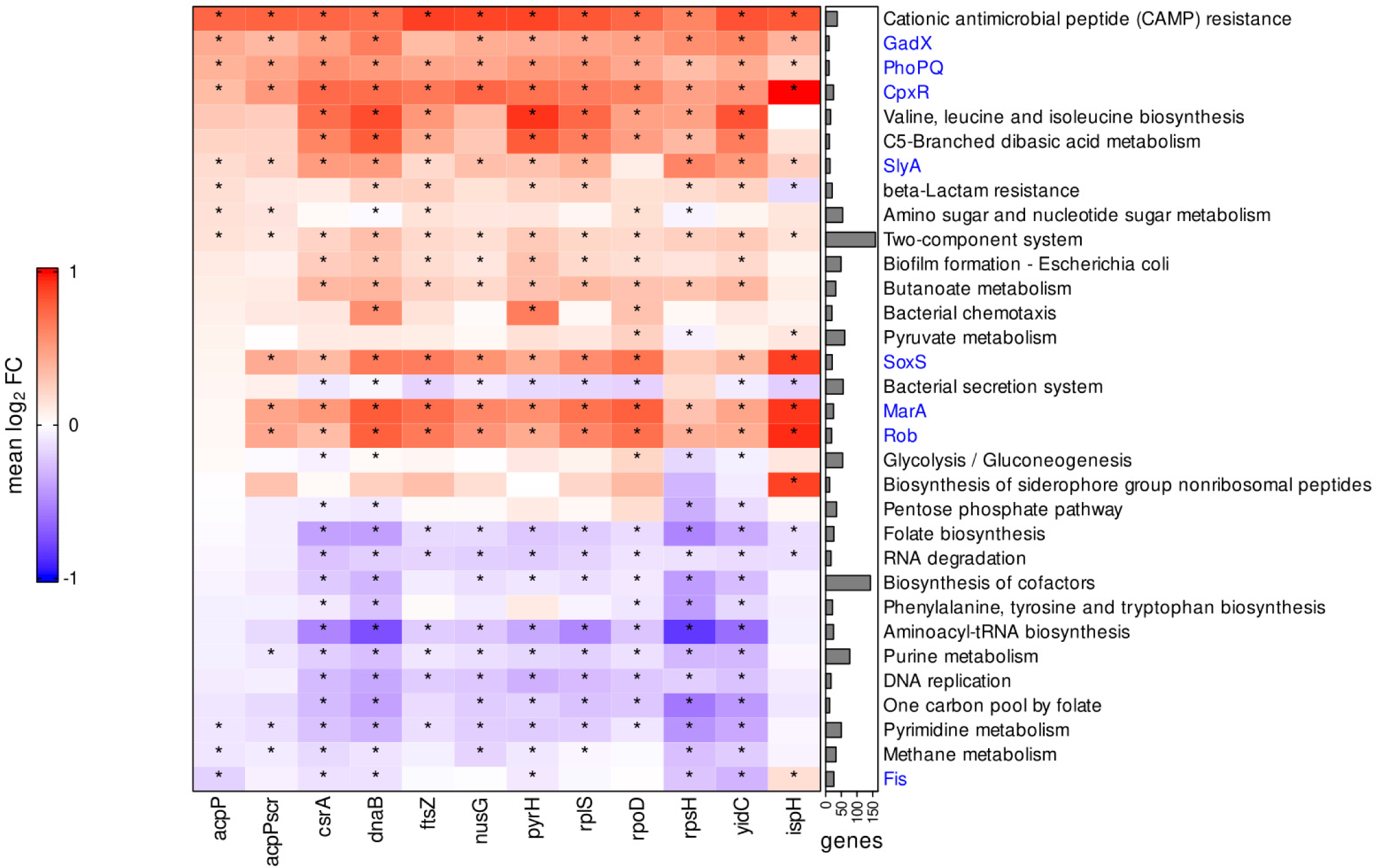
Gene set enrichment analysis. Global enrichment analysis of differentially expressed genes classified according to annotated KEGG pathways and manually added regulons (marked in blue, retrieved from RegulonDB annotations for *E. coli* K-12). Enrichment analysis shows a common induction of cationic antimicrobial peptide (CAMP) resistance, GadX, PhoPQ, CpxR, and Two-component system regulons. Asterisks (*) indicate statistically significant (*P*-value adj < 0.05) gene sets. Right bar chart shows the number of genes included within the single KEGG pathways/regulons. Up- (red) and downregulated (blue) pathways are indicated as median log_2_ fold change (FC).

We then analyzed the regulation of the direct PPNA targets in each condition (Figure 8, “targets”). In addition to (KFF)_3_K-*acpP* (4.8-fold change), only 4 out of the 10 tested PPNAs triggered a substantial reduction of their target mRNA level, namely *dnaB* (2.5-fold change), *pyrH* (4.9-fold change), *rpoD* (8.2-fold change), and *yidC* (3.5-fold change). Notably, the latter three were not promising candidates based on MIC analysis (≥ 10 µM; Table 1). Most other PPNAs showed moderate but non-significant reduction of their respective target, except (KFF)_3_K-*rpsH* which did not appear to impact *rpsH* mRNA levels, despite blocking *in vitro* translation of its target (Figure 5B) and inhibiting growth of UPEC 536 (Table 1). Importantly, the scrambled control did not mediate mRNA depletion for any of the essential gene targets. Based on these findings we conclude that rapid PPNA-triggered reduction of target transcript levels is not a universal feature of PPNA antisense inhibition and is also not associated with MIC (Table 1).

**Figure 8:**
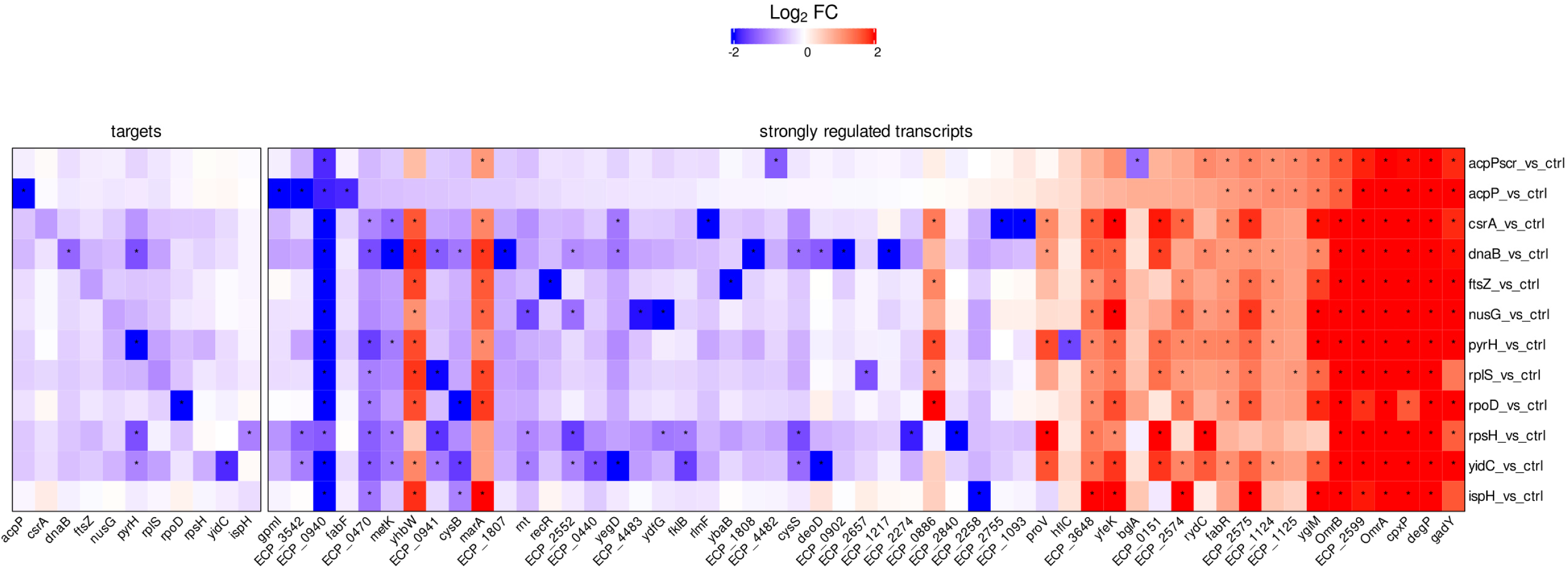
Transcriptomic profiling shows that substantial PPNA-triggered target mRNA depletion is not a universal mechanism, but reveals the induction of a common PNA sequence-independent response. UPEC 536 cells (10^6^ cfu/ml) were challenged with each of the PPNA conjugates indicated on the right (5 µM), or an equal volume of water as control. Cells were harvested 15 min post treatment for RNA isolation and subsequent RNA-seq analyses. Besides each PPNA’s target gene (left panel), top-regulated transcripts for each condition were included into the heatmap (right panel). Differential expression (log_2_ fold changes (FC), red and blue color code denotes up- and down-regulation, respectively) of each gene is shown relative to the untreated control. Asterisks (*) indicate significantly regulated genes (absolute FC >2) with a false discovery rate (FDR) adjusted *P*-value < 0.001. Comparisons for all conditions include triplicate RNA-seq samples.

### Decoding direct and indirect PPNA-triggered transcriptomic responses

Finally, we extended the analysis to the five top up- and downregulated transcripts for each condition, shown in one heatmap (Figure 8, “strongly regulated transcripts”). Closer inspection of the gold standard (KFF)_3_K-*acpP* PNA, which has been recently employed to analyze transcriptomic changes of PPNA-treated *Salmonella* (26), revealed a limited transcriptomic response in UPEC (Figure 8; Dataset S1). While 43 genes were significantly induced, only 16 genes were significantly reduced upon *acpP* PNA treatment. Specifically, in addition to *acpP* itself, (KFF)_3_K-*acpP* triggered pronounced depletion of *fabF* (3.5-fold change), a gene co-transcribed with *acpP.* (KFF)_3_K-*acpP* also specifically depleted *gpmI* (2,3-bisphosphoglycerate-independent phosphoglycerate mutase; 5.5-fold change) and ECP_3542 (glycerophosphoryl diester phosphodiesterase; 4.7-fold change). This is likely due to the presence of a fully complementary 9 base PNA binding site within the translation initiation sites of *gpmI*. Similarly, two transcripts, ECP_4482 and *bglA,* were exclusively depleted by the (KFF)_3_K-*acpP*-scrambled PNA and both have 8 base complementary binding sites in proximity to their respective annotated translational start sites (Figure 8). These observations suggest that an 8- to 9mer PNA can be sufficient to trigger depletion of a transcript.

PPNAs also elicited a common and thus sequence-independent regulation of certain transcripts. We identified *ompF* (outer membrane protein F, ECP_0940), one of the most abundant porins in the outer membrane (OM) of *E. coli* (56), among the mRNAs depleted in all conditions (approximately 3.6-fold change). This suggests that *ompF* is part of a global and likely indirect transcriptional response to PPNA exposure of UPEC. The strongest increase in expression among all conditions was found for the three small RNAs (sRNAs) GadY, OmrA and OmrB, as well as the mRNAs *degP, cpxP*, and ECP_2599 (RaiA). The sRNAs OmrA and OmrB regulate OM protein composition (57–59), and may be indicative of global envelope remodeling upon PPNA exposure. The transcript *degP*, also known as *htrA*, encodes a periplasmic serine endoprotease involved in protein quality control and stress response that has both chaperone and protease activities (60, 61). CpxP – regulated by the CpxA/R inner membrane stress response – is a periplasmic adaptor protein and the negative feedback regulator of CpxA, which fine tunes the CpxA/R pathway (62–64). Further, CpxP acts as a scaffold protein for delivery of misfolded periplasmic proteins for DegP-mediated proteolysis (65, 66). We also observed significant upregulation of ECP_2575, encoding the extracytoplasmic function sigma-E factor (σE), a master regulator of the OM stress response in Gram-negative bacteria (64, 67).

Taken together, our findings demonstrate that treatment of UPEC 536 with different PPNA constructs elicits a common envelope stress response, driven by σE and Cpx, and very likely triggers remodeling of the bacterial membrane, as indicated by depletion of *ompF* and induction of OmrA/B.

### The growth inhibitory capacity of a PNA is affected by its length

It has been experimentally shown that 9-11mer PNAs have the highest efficacy in *E. coli* (6–8). The optimal length was proposed to be 10 nucleobases, since longer oligomers possess suboptimal uptake properties while shorter ones show reduced duplex stability (7). However, this conclusion is almost exclusively based on the analysis of PNAs that target *lacZ* or *acpP* (6–8). We therefore set out to comprehensively analyze PNA length effects and to elucidate whether the proposed optimal length of a 10mer PNA has general validity (7) (Table 1). To this aim, we systematically analyzed the growth inhibitory capacity of 8mer, 9mer, and 11mer PNAs, using the same assay conditions as described for their 10mer counterparts.

Surprisingly, MIC assays using 9mer PNAs targeting each of the essential genes listed in Table 1 revealed that almost all 9mer PPNAs caused a similar, in some cases even stronger, growth inhibition compared to their 10mer counterparts (Table 1; Figures S1 and S3). Specifically, 9mer PPNAs targeting *dnaB, ftsZ,* or *rpsH* led to slightly improved growth inhibition compared to 10mers. To determine if the PNA length could be reduced even further, we tested 8mer PPNAs for the most efficient PNA targets, *i.e. dnaB, ftsZ, rpsH*, and *acpP.* Three out of four constructs showed a 2-4 fold higher MIC than the respective 9mer PPNA (Table 1; Figure S4), indicating that 9mer, but not 8mer, PPNAs can be more efficient in mediating growth inhibition compared to their 10mer counterparts. This might be attributed to either improved uptake or to increased target specificity, due to less frequent off-targeting (8).

To evaluate whether increased binding affinity improves growth inhibition (8), we also tested extended 11mer PPNAs in MIC assays (Table 1; Figure S5). We observed similar or less efficient growth inhibition for the majority of tested 11mer PPNAs compared to 10mer molecules (Table 1). One notable exception is (KFF)_3_K-*acpP*, which had an identical MIC regardless of length. Overall, we conclude based on these data that 9mer PNA molecules can be at least as efficient in mediating growth inhibition as their 10mer counterparts (Table 1).

## DISCUSSION

Our current mechanistic understanding of PNA function is mostly derived from studies involving a small number of PNA targets (1,6–8,13,18,26,29,30,68). Here, we present a systematic investigation of the antibacterial activity and mode of action of eleven PPNAs targeting different essential genes in UPEC. Our analysis revealed a broad distribution of growth inhibitory capacities. Below, we will discuss specific features of PPNA activity and PPNA design that might impact their antibacterial efficacy.

### Aspects of PNA activity and implications for PNA design

Antisense PNAs targeting essential bacterial genes have been shown to possess antibacterial activity via selective inhibition of their targets (reviewed elsewhere (4, 5)). The most commonly studied PNA target is *acpP* (5, 8), which is strongly conserved and one of the most highly expressed genes in UPEC (Figures 1B and 5A). Importantly, *acpP* emerged as the most vulnerable essential target gene in our systematic analysis, based on the level of growth inhibition upon *acpP* targeting. What makes then *acpP* PNA so potent? One obvious explanation could be that the PNA used here also depletes the mRNAs of three additional genes, *i.e., fabF*, *gpmI* and ECP_3542 (*ugpQ*), as part of its off-target activity. However, to the best of our knowledge, neither of the encoded proteins, *i.e.,* 3-oxoacyl-[acyl carrier protein] synthase 2, 2,3-bisphosphoglycerate-independent phosphoglycerate mutase and glycerophosphoryl diester phosphodiesterase, respectively, is essential for the growth of *E. coli* (54, 69). Further, the lethal phenotype triggered by *acpP* PNA treatment in *E. coli* can be rescued by a plasmid-encoded *acpP* allele designed to have low complementarity to *acpP* PNA, confirming that depletion of *acpP* causes growth inhibition (8).

Another explanation might be the strong vulnerability of bacterial cells to *acpP* depletion, referred to as gene stringency, which exceeds that of other essential genes in *E. coli* (29). In particular, a 40 % reduction in *acpP* mRNA levels leads to a 50 % growth inhibition, whereas less stringency was demonstrated for *ftsZ, murA* and *fabI* (29). The vulnerability of *acpP* is further supported by a recent CRISPR interference (CRISPRi) study in *Mycobacterium tuberculosis*, which demonstrated that even incomplete inhibition of particular essential genes can severely affect bacterial fitness, even killing, while the bacteria are less sensitive to the depletion of other essential genes (70). The fatty acid biosynthesis pathway was highly enriched for vulnerable genes, including the *acpP* ortholog *acpM*, in *Mycobacterium tuberculosis* (https://pebble.rockefeller.edu/; (70)), suggesting that this trait could be deeply conserved across the bacterial phylogeny. Similarly, a recent CRISPRi study in *E. coli* also found differential vulnerability to transcript depletion among essential genes (71). Follow-up studies on the full essential gene targetome would be needed to classify particularly vulnerable pathways and genes for PNA treatment and to identify the most attractive targets, similar to the CRISPRi studies performed in *Mycobacterium* (70) or *E. coli* (71). Such analyses would also clarify whether the connectedness of a target protein’s interaction network is a universal predictor of the growth inhibitory activity of a PNA, as recently suggested (19).

Designed to target the translational start site, the mechanism of action of PNAs is generally ascribed to the inhibition of target mRNA translation (6,8,13). We measured the effects of PNAs on mRNA translation using an *in vitro* translation system, which has been applied to monitor the translation rate of a target mRNA in the presence of its antisense sRNA (72–76). Previously, a similar approach was adopted to verify PNA-mediated translational inhibition of a *Rickettsia*-encoded gene (77). The *in vitro* translation system constitutes a straightforward method to predict the efficacy of PNA-mediated inhibition of translation *in vitro*. Additionally, it enables the direct analysis of PNA-mediated translational inhibition independent of *in cellulo* target mRNA stability. However, this system may not fully reconstitute intracellular target mRNA conformation or accessibility. In addition, the *in vitro* system is independent of the PNA uptake efficiency into the bacterial cell, which is a critical step that contributes to the activity of a PPNA. These considerations might explain why the efficacy of inhibiting *in vitro* translation is not directly linked to PNA-mediated growth inhibition. In particular, the absence of an *in vitro* effect on RplS translation is puzzling, given the PNA-mediated growth reduction at high concentrations (Figure 5B, Table 1), though this may reflect a higher intracellular molar ratio than tested in our *in vitro* translation assay. Besides, it is also conceivable that the growth inhibition mediated by the 10mer *rplS* PNA at ≥ 10 µM (Table 1) is due to the off-target depletion of ECP_2657 (Figure 8), which encodes the essential protein alanyl-tRNA synthetase. While this possibility remains to be experimentally tested, the complementarity of the *rplS* PNA to 9 nucleotides within the translation initiation site of ECP_2657 makes this a likely scenario. An alternative strategy to monitor PNA-mediated effects on translation *in vivo* is ribosome profiling (Ribo-seq) (78), but the establishment of this approach will require further optimization to minimize the starting material needed (79) due to the current costs of synthesizing PNA molecules.

Apart from their predicted mode of action, *i.e.* sterically blocking ribosome binding and thus inhibiting translation, several studies in *E. coli* and other gram-negative bacteria have shown that PNAs concomitantly trigger target mRNA depletion, including PNAs targeting *acpP, fabI, ftsZ, murA*, and *rpoD* (18,26,27,29–31). Since we recently demonstrated rapid PPNA-triggered depletion of *acpP* in *Salmonella* within 5 min of exposure (26), we reasoned that target mRNA depletion might be a universal rapid response caused by translational inhibition. However, based on our RNA-seq data a significant depletion of the PNA target is not universally observed after 15 min PPNA-exposure in UPEC (Figure 8) and is not associated with the PPNA’s growth inhibitory capacity (Table 1). Based on previous genome-wide studies of mRNA half-lives and translation rates in *E. coli*, we exclude a strong contribution of target mRNA stability (80) and translation efficiency (28) in PPNA-mediated depletion (Figure 8). It thus remains unclear what factors might determine the degree of target mRNA depletion upon PPNA treatment or whether some target mRNAs might have a delayed decay. Nevertheless, our data show that rapid target mRNA depletion is not a universal consequence of PPNA treatment.

Previous reports suggested that 10mer PNAs confer maximal activity, at least when targeting *E. coli acpP* (7) or *lacZ* (8). This length optimum reflects a balance of (i) PNA binding affinity, (ii) PPNA uptake efficiency, and (iii) the probability of off-target binding (7,8,32–34). Unexpectedly, our analysis revealed that all but one 9mer PNAs (*i.e., rplS* PNA) possess a similar or even superior ability to inhibit growth compared to 10mers (Table 1). Therefore, generalizations on an optimal PNA length based on one individual gene are not universally valid. Curiously, generation of a 9mer PNA by truncating the commonly used 10mer *acpP* PNA on the N-terminus dampens its antibacterial activity against *E. coli* K-12 (7), while truncation of the same 10mer *acpP* PNA on its C-terminus does not affect the growth inhibitory capacity of the resulting construct in UPEC (Table 1). Whether these 9mer *acpP* PNAs would have different antibacterial activities if tested in the same organism is unclear. Nevertheless, our results demonstrate that using 9mer instead of 10mer PNAs can improve antibacterial activity, although this may require optimization of the truncated PNA sequence. A one-nucleobase difference might seem incremental but we expect 9mer PNAs to have fewer off-targets, which will be important for PNA applications seeking to rewire bacterial gene expression with high specificity.

The stability of a PNA/RNA duplex, measured by its melting temperature, has been proposed to affect PNA efficacy (7,8,32–34). Specifically, a positive correlation of the melting temperature and growth inhibitory capacity has been demonstrated for 8-10mer *acpP* PNAs (7). However, our systematic analysis of 8-10mer PNAs targeting 11 essential genes does not support a direct correlation between the predicted melting temperature and the growth inhibitory capacity (Figure S6), although we find that the predicted melting temperature is associated with the capacity of a PNA to inhibit translation of its target *in vitro*. This suggests that other factors dictate the antisense efficacy and thus antibacterial activity of a PNA *in cellulo*. While target mRNA secondary structure has been assumed to play a key role, it has recently been excluded as a crucial determinant of a PNA’s growth inhibitory activity (19).

Apart from targeting essential genes, PNAs can also be applied to reduce the expression of antibiotic resistance genes and thereby to resensitize (multi)drug resistant bacteria to antibiotics (81–83). In addition, combined administration of PNAs targeting essential but also non-essential genes together with conventional antibiotics has been shown to increase the sensitivity of multidrug resistant bacteria to the respective compound (19). These types of combination treatment may be especially important to combat UPEC, due to the frequent emergence of drug resistant clinical isolates (23, 24).

### Defining the PPNA-triggered transcriptomic landscape in UPEC

PNAs conjugated to (KFF)_3_K peptide, which were used throughout this study, are around 4 kDa in size and are thus substantially larger than conventional antibiotics (84, 85). In gram-negative bacteria, uptake of several cationic antibiotics, such as kanamycin, beta-lactams or quinolones, is facilitated by passive diffusion through OM porins, such as OmpF (86–92). In contrast, (KFF)_3_K-coupled PNAs follow a different mechanism. Specifically, positively charged (KFF)_3_K-PNA molecules bind to and destabilize the negatively charged LPS layer of the OM of gram-negative bacteria to facilitate their periplasmic translocation (93, 94). A yet unknown peptidase partially cleaves (KFF)_3_K in the extracellular and periplasmic space (94). The truncated PPNA molecules can then cross the inner membrane via the SbmA transporter in *E. coli* (27, 94). Curiously, instead of a depletion of *sbmA* transcript levels to promote resistance, our RNA-seq data revealed increased mRNA levels of *sbmA* in all but one of the analyzed PPNA conditions. It is tempting to speculate that this is caused by a strong induction of σE and Cpx stress responses (95, 96), which will be discussed in more detail below.

Following SbmA-mediated uptake, PNAs are released into the cytoplasm and can exert their function as antisense drugs (27, 94). Due to their size and capacity for membrane permeabilization (97), it is not surprising that PPNAs induced a set of genes implicated in two well-known envelope stress pathways. Specifically, we found strong upregulation of members of the σE and Cpx responses (Figures 7 and 8), which, together with the Rcs (regulator of capsule synthesis), Bae (bacterial adaptive response), and Psp (phage shock protein) responses, form protective signal transduction cascades that ameliorate the detrimental effects of specific stressors (reviewed in (64)). While the σE response is triggered upon OM stress, usually via sensing of misfolded OMPs or aberrant forms of LPS, the Cpx response is related to inner membrane stress (64,98,99). Since excessive activation of envelope stress pathways can have toxic effects on the cell, their activation is balanced via negative feedback regulators (64). In particular, RseA (ECP_2574) and CpxP counterbalance activation of the σE and Cpx response, respectively, and are part of the respective stress response regulons (96,100–102). PPNA-induced transcript levels of *rseA*, *cpxP* and *degP*, which encodes a periplasmic protease that is transcriptionally activated by both σE and Cpx, reflect at least a partial activation of both pathways (Figure 8) (95,96,103–106). Thus, UPEC rapidly reacts to PPNA treatment via outer and inner membrane stress response pathways.

Recently, constitutive induction of the Cpx response has been linked to the development of tolerance to (RXR)_4_XB-coupled PNAs in *E. coli*. However, efficacy was unaffected under the tested conditions if the PNAs were coupled to (KKF)_3_K instead (51). In addition, aberrant activation of σE in *E. coli* has been shown not to confer tolerance to antimicrobial peptides (AMPs), such as arginine-rich CPPs coupled to PNAs, but there is lack of information regarding the effect of σE on (KFF)_3_K-PNA activity (51, 107). Therefore, it remains to be seen whether induction of the Cpx and σE stress response pathways can confer some degree of tolerance or resistance to PPNAs in UPEC.

We also observed universal PPNA-mediated depletion of the *ompF* mRNA (Figure 8), very likely due to its direct regulation by both the σE and Cpx stress responses (96,108–110). The Cpx response is also known to indirectly dampen the expression of *ompF.* In particular, Cpx induces the expression of *marA*, which in turn triggers the expression of the *ompF*-targeting sRNA MicF (58,111–115). Both *marA* and MicF were found to be highly induced in most of the PPNA conditions in our data sets (Figures 8 and S7). However, whether *ompF* mRNA depletion has physiological consequences or may influence PPNA uptake remains uncertain.

Overall, our study reveals the efficacy of PPNAs against UPEC and presents a systematic analysis of eleven PNAs targeting different essential genes. We observe strong antibacterial effects for certain 10mer PPNAs independent of target gene abundance and provide evidence that 9mer PNAs, instead of their commonly used 10mer counterparts, can alternatively be used to inhibit bacterial growth. Global RNA-seq analysis indicate that rapid PPNA-mediated target mRNA depletion is not universal, but that PPNA treatment induces a common set of envelope stress responses and membrane remodeling factors. Further mechanistic studies will be needed to determine whether these responses may influence resistance to or tolerance of PPNAs. Nonetheless, our data will guide future PPNA design and help to better understand the physiological changes triggered by PPNA exposure.

## DATA AVAILABILITY

Our RNA-seq data set has been uploaded on repositories and is available upon request.

## Supporting information

Supplementary Data Set S1

## ACKNOWLEDGEMENT

We thank Barbara Plaschke for extensive technical assistance and Tobias Kerrinnes for laboratory organization. We thank Regan Hayward for computational analysis in the early phase of this project. We especially thank Anke Sparmann for helpful comments and edits on the manuscript. We further thank Tina Lence and Svetlana Ðurica-Mitić for constructive comments on the manuscript. We also thank the Core Unit Systems Medicine (Julius-Maximilians-University, Würzburg) for handling our RNA samples for sequencing.

