## Supplementary Material for "Comprehensive analysis of PNA-based antisense antibiotics targeting various essential genes in uropathogenic *Escherichia coli*"

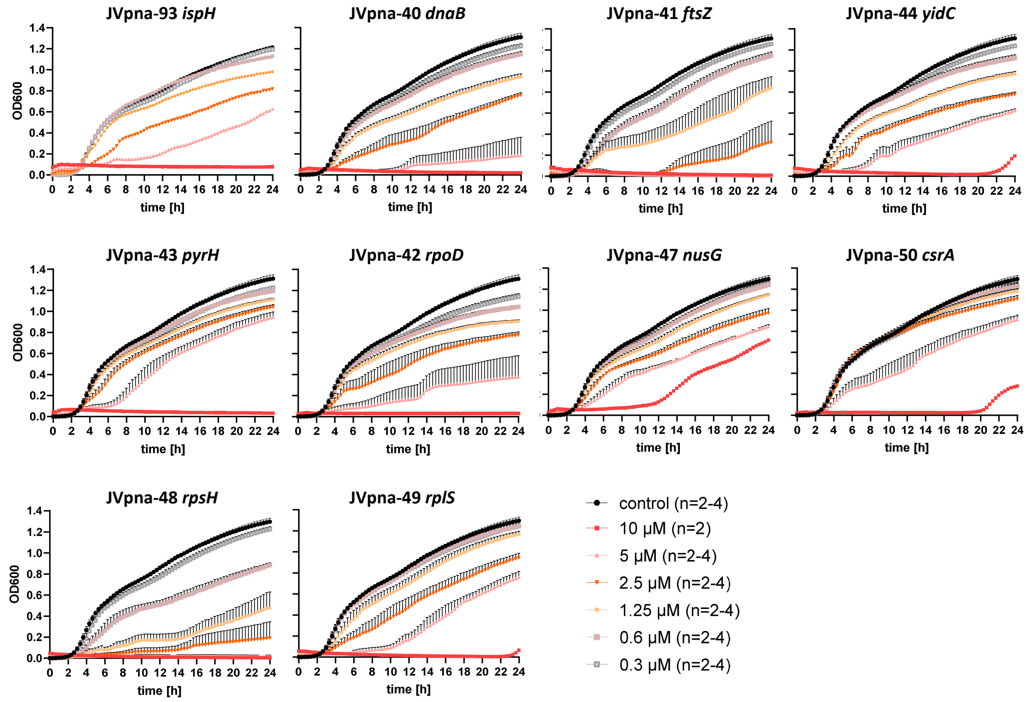


**Fig. S1. Analyzing the growth inhibitory capacity of 10mer PPNAs targeting selected low to high abundant essential genes.** UPEC 536 cells ($10^5$ cfu/ml) were exposed to serial dilutions ranging from 10 µM to 0.3 µM of $(KFF)_3K$-coupled 10mer PPNAs, or water as control (black curves), and growth was monitored within 24 h. Minimum inhibitory concentrations (MICs) were determined for the applied PPNAs against UPEC 536 and are summarized in Table 1. Growth is shown as $OD_{600}$ (y-axis) over time (indicated in hours, x-axis).The experiments were performed two to four times. Curves represent the mean of all biological replicates and error bars indicate standard error of the mean.

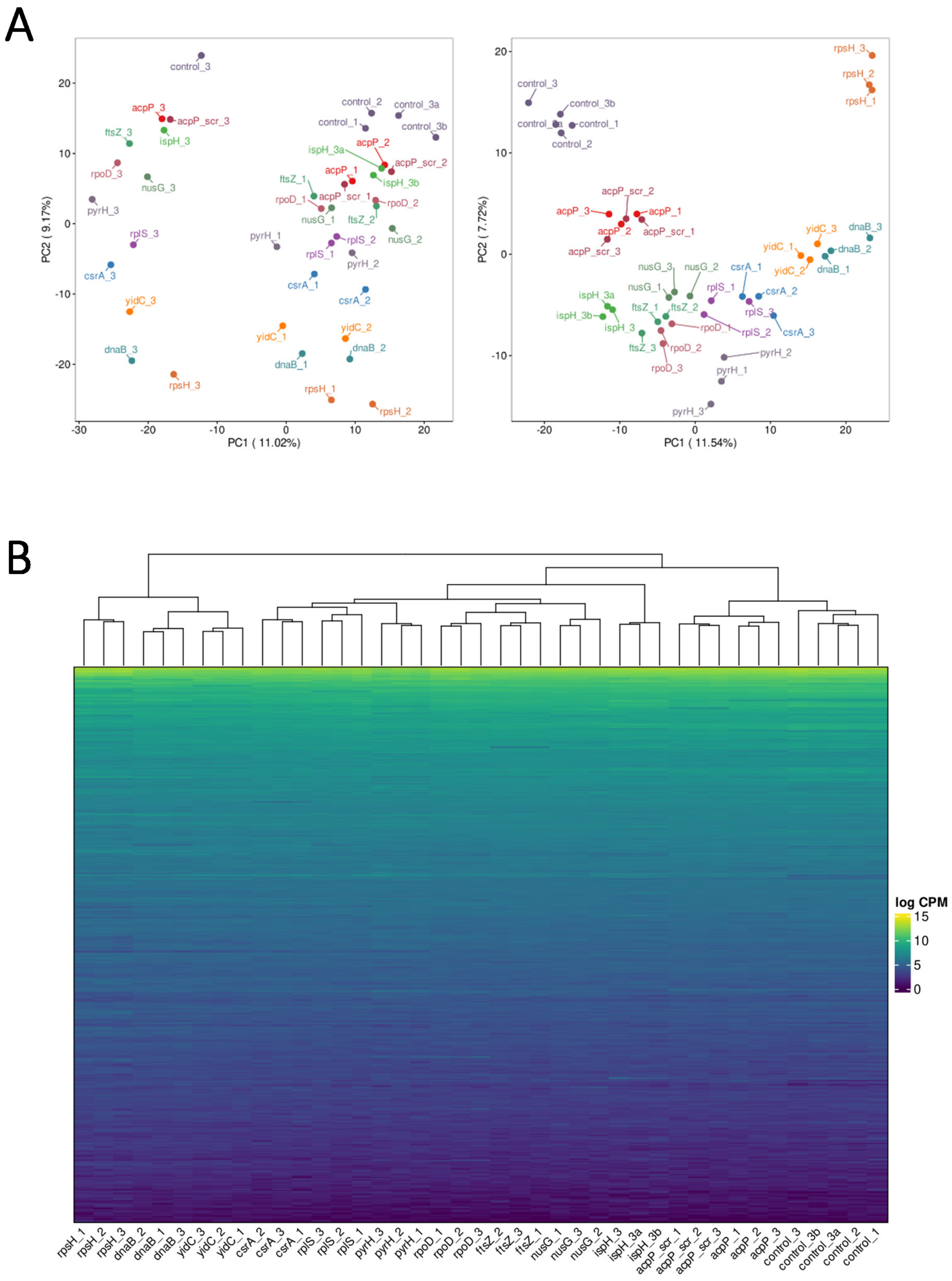


**Fig. S2. Transcriptomic profiling of UPEC 536 upon short-term exposure to various essential gene-targeting PPNAs.** UPEC 536 cells ($10^6$ cfu/ml) were exposed to 5 µM of $(KFF)_3K$-coupled 10mer PPNAs targeting various essential genes. Scrambled PPNA was applied as sequence-unrelated control. An equal volume of water was used as untreated control. After 15 min, total RNA was extracted and subjected to RNA-seq analyses. **(A)** Principal component analysis (PCA) of all 12 PPNA conditions (n=3) and the untreated controls (n=5) was performed. Left and right panels show PCA before and after batch effect removal, respectively. The different treatment conditions separate into four rough clusters, as shown in the PCA blot after batch removal (right panel). **(B)** Hierarchical cluster analysis (HCA) was performed after batch effect removal. Abundance of all detected transcripts are depicted as log CPM (counts per million). All biological replicates per treatment condition cluster together.

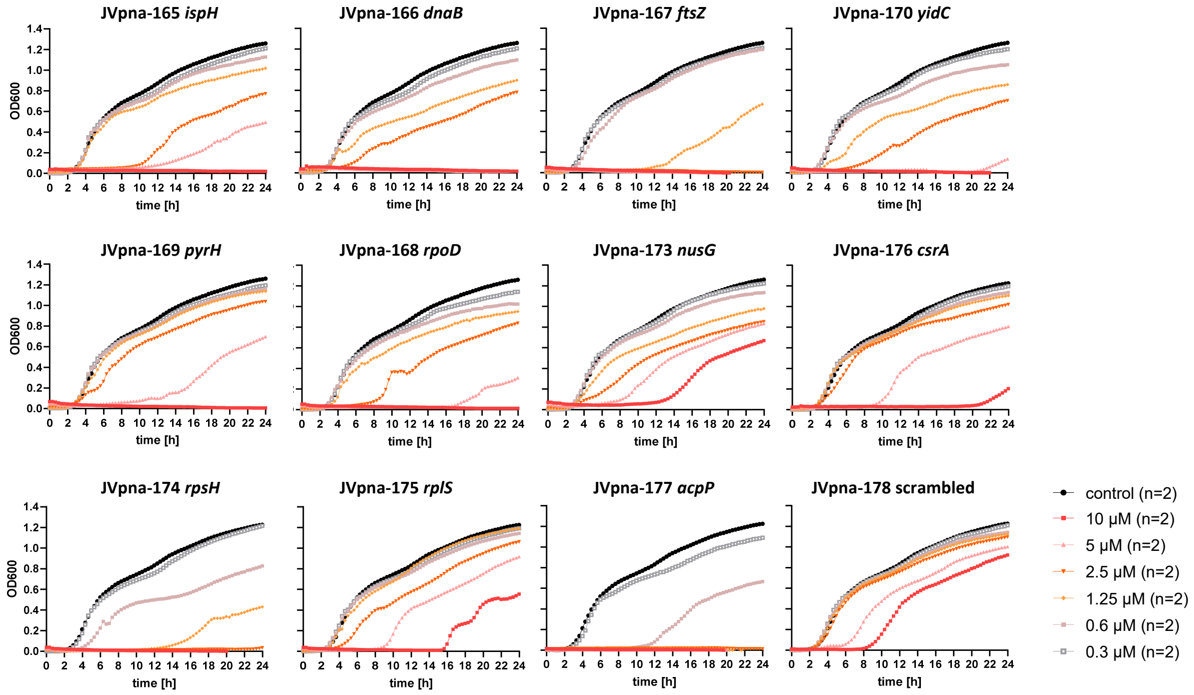


**Fig. S3. Analyzing the growth inhibitory capacity of 9mer PPNAs targeting selected low to high abundant essential genes.** UPEC 536 cells ($10^5$ cfu/ml) were exposed to serial dilutions ranging from 10 µM to 0.3 µM of $(KFF)_3K$-coupled 9mer PPNAs, or water as control (black curves), and growth was monitored within 24 h. Minimum inhibitory concentrations (MICs) were determined for the applied PPNAs against UPEC 536 and are summarized in Table 1. Growth is shown as $OD_{600}$ (y-axis) over time (indicated in hours, x-axis).The experiment was performed two times and curves represent the mean.

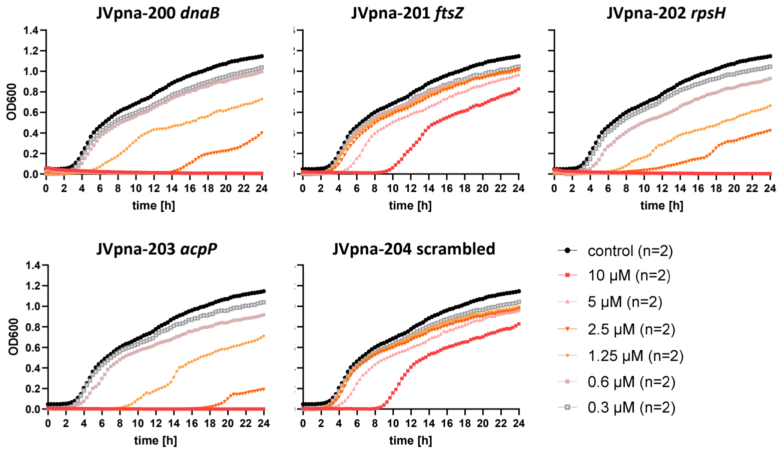


**Fig. S4. Analyzing the growth inhibitory capacity of 8mer PPNAs targeting selected low to high abundant essential genes.** UPEC 536 cells ($10^5$ cfu/ml) were exposed to serial dilutions ranging from 10 µM to 0.3 µM of $(KFF)_3K$-coupled 8mer PPNAs, or water as control (black curves), and growth was monitored within 24 h. Minimum inhibitory concentrations (MICs) were determined for the applied PPNAs against UPEC 536 and are summarized in Table 1. Growth is shown as $OD_{600}$ (y-axis) over time (indicated in hours, x-axis). The experiment was performed two times and curves represent the mean.

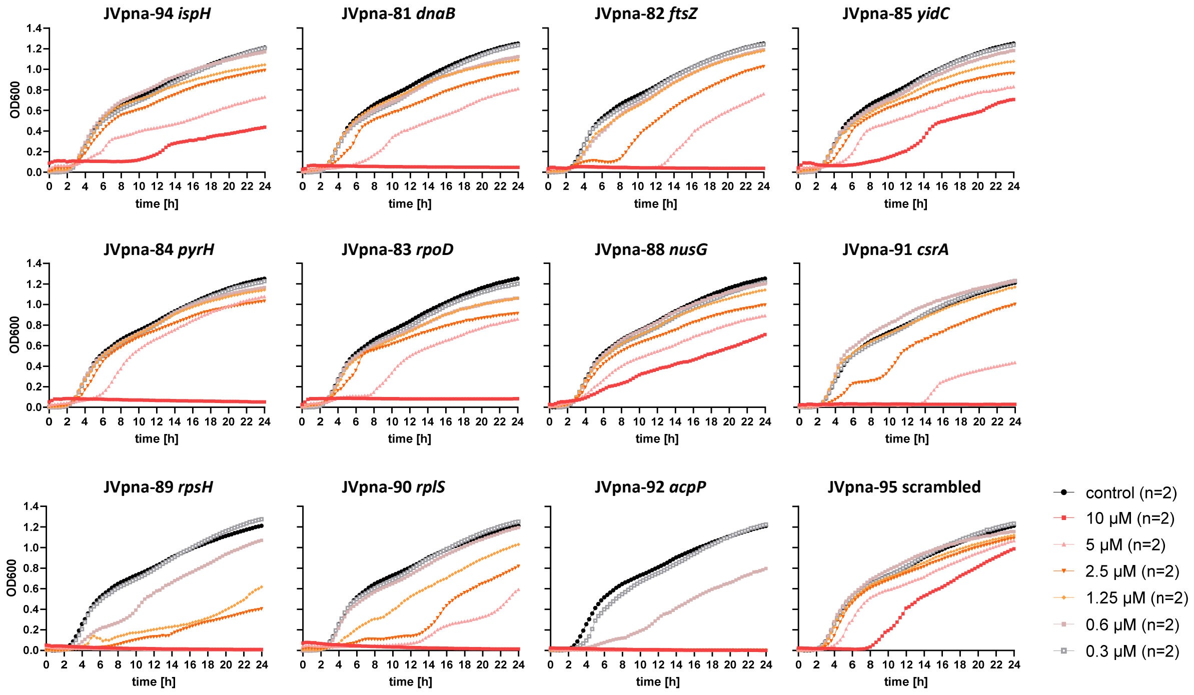


**Fig. S5. Analyzing the growth inhibitory capacity of 11mer PPNAs targeting selected low to high abundant essential genes.** UPEC 536 cells ($10^5$ cfu/ml) were exposed to serial dilutions ranging from 10 µM to 0.3 µM of (KFF)$_3$K-coupled 11mer PPNAs, or water as control (black curves), and growth was monitored within 24 h. Minimum inhibitory concentrations (MICs) were determined for the applied PPNAs against UPEC 536 and are summarized in Table 1. Growth is shown as $OD_{600}$ (y-axis) over time (indicated in hours, x-axis). The experiment was performed two times and curves represent the mean.

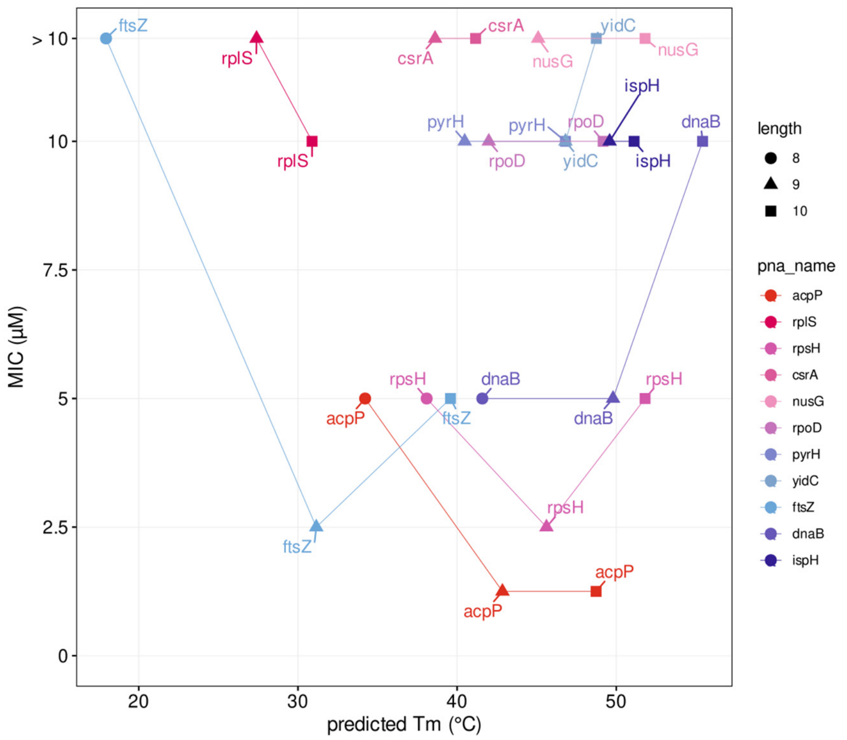


**Fig. S6.The antibacterial activity (MIC) of a PPNA is not associated with its predicted melting temperature.** MICs (shown in µM, y-axis) were derived from Table 1 (if the MIC lies between two concentrations, the higher one was chosen for creating this plot). Melting temperatures (Tm, shown in °C, x-axis) were predicted as described in the materials and methods section. This plot includes 8mer (●), 9mer (▲), and 10mer (■) essential gene targeting PPNAs (shown on the right). High to low target gene abundances are indicated from red to blue coloring, according to Figure 5.

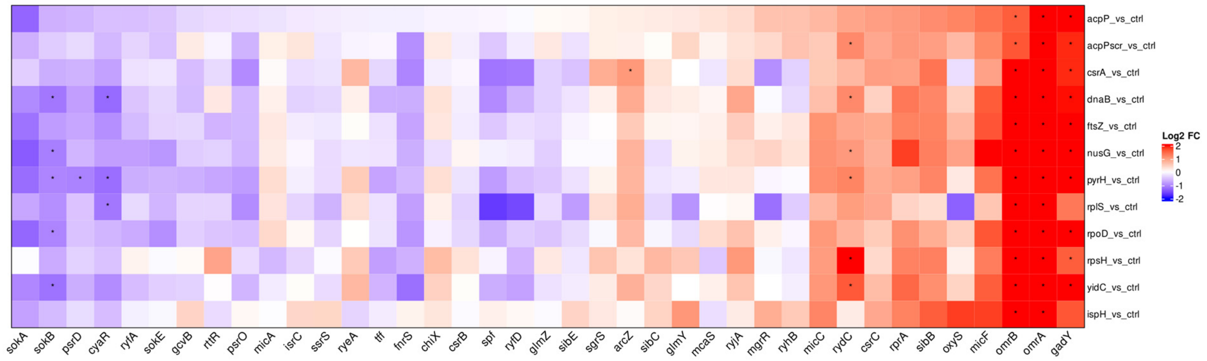


**Fig. S7. sRNA expression profiles upon short-term exposure to various essential gene-targeting PPNAs in UPEC 536.** Experimental setup is described in Fig. S2. Comparison of the RNA-seq dataset for the tested PPNA constructs versus control (as indicated on the right side) includes all biological replicates for each condition. Detected sRNAs are indicated on the bottom. Color code denotes $\log_2$ fold change (FC), with red to blue indicating up- to down-regulation, respectively. sRNA with an absolute fold change >2 (down-regulated log2 < –1, up-regulated log2 > 1) and an false discovery rate (FDR) adjusted P-value < 0.001 are considered as significantly differentially regulated and are marked with an asterisk (*).

**Table S1**. **Oligonucleotide sequences for generation of *gfp* fusion constructs.** Table includes internal oligonucleotide number (JVO-), target gene name, locus tag (ECP_) or PCR template for *gfp* amplification (pXG-10) (1), and the respective oligonucleotide sequence in 5’ to 3’ orientation. Fw: forward, rv: reverse. Underlined sequence: T7 promoter sequence, italic sequence: *gfp* overlap.

| Oligo (JVO-) | Gene | Locus tag (ECP_) | Sequence (5’ to 3’) |
|---|---|---|---|
| 19738 | *ispH* - fw | 0027 | TAATACGACTCACTATAGATTGAAGTGCTGGAAATCGA |
| 19739 | *ispH* - rv | | *TCCAGTGAAAAGTTCTTCTCCTTTGCTAGC*GCGGTCTACCCCGGCACAAAA |
| 19736 | *dnaB* - fw | 4268 | TAATACGACTCACTATAGAAGTTTCGACCCATTCTTTA |
| 19737 | *dnaB* - rv | | *TCCAGTGAAAAGTTCTTCTCCTTTGCTAGC*GCGTTCGCGGGGTTCAGCCTG |
| 19740 | *ftsZ* - fw | 0097 | TAATACGACTCACTATAGATTACGGCCTCAGGCGACAG |
| 19741 | *ftsZ* - rv | | *TCCAGTGAAAAGTTCTTCTCCTTTGCTAGC*GCCGATGACTTTAATCACCGC |
| 19746 | *yidC* - fw | 3906 | TAATACGACTCACTATAGCCGTCCCGCCCGGACCATTT |
| 19747 | *yidC* - rv | | *TCCAGTGAAAAGTTCTTCTCCTTTGCTAGC*GAAAGACACGAACAGCAAAGC |
| 19744 | *pyrH* - fw | 0179 | TAATACGACTCACTATAGATTAATTCATTTCAATCGTT |
| 19745 | *pyrH* - rv | | *TCCAGTGAAAAGTTCTTCTCCTTTGCTAGC*ACTCAACTTAAGCAGAATGCG |
| 19742 | *rpoD* - fw | 3157 | TAATACGACTCACTATAGCCCAACACCAACCTCATGAA |
| 19743 | *rpoD* - rv | | *TCCAGTGAAAAGTTCTTCTCCTTTGCTAGC*CTTACCACGGGTGACAAGAAG |
| 19752 | *nusG* - fw | 4195 | TAATACGACTCACTATAGGTTCGCCTGGTATCCTTTAT |
| 19753 | *nusG* - rv | | *TCCAGTGAAAAGTTCTTCTCCTTTGCTAGC*ACCGGAAAACGCCTGAACGAC |
| 19758 | *csrA* - fw | 2656 | TAATACGACTCACTATAGCAGAGAGACCCGACTCTTTT |
| 19759 | *csrA* - rv | | *TCCAGTGAAAAGTTCTTCTCCTTTGCTAGC*CTCATCCCCAATCATGAGGGT |
| 19754 | *rpsH* - fw | 3394 | TAATACGACTCACTATAGCTGGTAATTGTCACCAATTG |
| 19755 | *rpsH* - rv | | *TCCAGTGAAAAGTTCTTCTCCTTTGCTAGC*ACCGTTACGGATACGGGTCAG |
| 19756 | *rplS* - fw | 2607 | TAATACGACTCACTATAGCCCCCGAGATATCAGTTTAC |
| 19757 | *rplS* - rv | | *TCCAGTGAAAAGTTCTTCTCCTTTGCTAGC*TACGTCCTGCTTCATCTGCTC |
| 19760 | *acpP* - fw | 1086 | TAATACGACTCACTATAGAACCATCGCGAAAGCGAGTT |
| 19761 | *acpP* - rv | | *TCCAGTGAAAAGTTCTTCTCCTTTGCTAGC*GCCCAGCTGTTCGCCGATAAT |
| **Oligo (JVO-)** | **Gene** | **Template** | **Sequence (5’ to 3’)** |
| 19762 | *gfp* - fw | pXG-10 | GCTAGCAAAGGAGAAGAACTTTTCAC |
| 19763 | *gfp* - rv | | TTATTTGTAGAGCTCATCCATGCC |

**Table S1: PPNA and peptide sequences.** Table includes internal PPNA and peptide number, target gene name, locus tag, peptide and respective PNA sequence (from N to C terminus orientation, PNA N to C terminus corresponds to nucleic acid 5' to 3' orientation), and the target region of the PNA relative to the translational start. X = 6-aminohexanoic acid; B = β-alanine

| PPNA# | gene | locus tag | Xmer | Peptide sequence (N to C) | PNA Sequence (N to C) | Target region |
|---|---|---|---|---|---|---|
| JVpna-21 | *acpP* | ECP_1086 | 10 | KFFKFFKFFK | CTCATACTCT | -5 to +5 |
| JVpna-22 | scrambled | - | | | TCACTATCTC | - |
| JVpna-18 | *acpP* | ECP_1086 | | RXRRXRRXRRXRXB | CTCATACTCT | -5 to +5 |
| JVpna-19 | scrambled | - | | | TCACTATCTC | - |
| JVpna-143 | *acpP* | ECP_1086 | | GRKKRRQRRRYK | CTCATACTCT | -5 to +5 |
| JVpna-97 | scrambled | - | | | TCACTATCTC | - |
| JVpna-196 | *acpP* | ECP_1086 | | Dap9 (2,3-Diaminopropionic Acid) | CTCATACTCT | -5 to +5 |
| JVpna-197 | scrambled | - | | | TCACTATCTC | - |
| JVpep-112 | - | - | - | KFFKFFKFFK | - | - |
| JVpep-113 | - | - | - | RXRRXRRXRRXRXB | - | - |
| JVpep-114 | - | - | - | GRKKRRQRRRYK | - | - |
| JVpna-200 | *dnaB* | ECP_4268 | 8 | KFFKFFKFFK | CTGCCATA | -1 to +7 |
| JVpna-201 | *ftsZ* | ECP_0097 | | | CAAACATA | -1 to +7 |
| JVpna-202 | *rpsH* | ECP_3394 | | | CTCATCTG | -3 to +5 |
| JVpna-203 | *acpP* | ECP_1086 | | | CTCATACT | -3 to +5 |
| JVpna-204 | scrambled | - | | | CACTATCT | - |
| JVpna-165 | *ispH* | ECP_0027 | 9 | KFFKFFKFFK | TCTGCATGT | -2 to +7 |
| JVpna-166 | *dnaB* | ECP_4268 | | | CTGCCATAG | -2 to +7 |
| JVpna-167 | *ftsZ* | ECP_0097 | | | CAAACATAG | -2 to +7 |
| JVpna-170 | *yidC* | ECP_3906 | | | TCCATCGTT | -4 to +5 |
| JVpna-169 | *pyrH* | ECP_0179 | | | CCATGTTTC | -5 to +4 |
| JVpna-168 | *rpoD* | ECP_3157 | | | CCATAAGAC | -5 to +4 |
| JVpna-173 | *nusG* | ECP_4195 | | | GACATCTCA | -4 to +5 |
| JVpna-176 | *csrA* | ECP_2656 | | | AGCATTCTT | -4 to +5 |
| JVpna-174 | *rpsH* | ECP_3394 | | | CTCATCTGT | -4 to +5 |
| JVpna-175 | *rplS* | ECP_2607 | | | CTCATAATT | -4 to +5 |
| JVpna-177 | *acpP* | ECP_1086 | | | CTCATACTC | -4 to +5 |
| JVpna-178 | scrambled | - | | | CACTATCTC | - |
| JVpna-93 | *ispH* | ECP_0027 | 10 | KFFKFFKFFK | TCTGCATGTT | -3 to +7 |
| JVpna-40 | *dnaB* | ECP_4268 | | | CTGCCATAGT | -3 to +7 |
| JVpna-41 | *ftsZ* | ECP_0097 | | | CAAACATAGT | -3 to +7 |
| JVpna-44 | *yidC* | ECP_3906 | | | TCCATCGTTA | -5 to +5 |
| JVpna-43 | *pyrH* | ECP_0179 | | | CCATGTTTCT | -6 to +4 |
| JVpna-42 | *rpoD* | ECP_3157 | | | CCATAAGACG | -6 to +4 |
| JVpna-47 | *nusG* | ECP_4195 | | | GACATCTCAG | -5 to +5 |
| JVpna-50 | *csrA* | ECP_2656 | | | AGCATTCTTT | -5 to +5 |
| JVpna-48 | *rpsH* | ECP_3394 | | | CTCATCTGTC | -5 to +5 |
| JVpna-49 | *rplS* | ECP_2607 | | | CTCATAATTT | -5 to +5 |
| JVpna-94 | *ispH* | ECP_0027 | 11 | KFFKFFKFFK | ATCTGCATGTT | -3 to +8 |

| | | | | | | |
|---|---|---|---|---|---|---|
| JVpna-81 | *dnaB* | ECP_4268 | | | CTGCCATAGTG | -4 to +7 |
| JVpna-82 | *ftsZ* | ECP_0097 | | | CAAACATAGTT | -4 to +7 |
| JVpna-85 | *yidC* | ECP_3906 | | | TCCATCGTTAG | -6 to +5 |
| JVpna-84 | *pyrH* | ECP_0179 | | | CCATGTTTCTT | -7 to +4 |
| JVpna-83 | *rpoD* | ECP_3157 | | | CCATAAGACGG | -7 to +4 |
| JVpna-88 | *nusG* | ECP_4195 | | | GACATCTCAGA | -6 to +5 |
| JVpna-91 | *csrA* | ECP_2656 | | | AGCATTCTTTG | -6 to +5 |
| JVpna-89 | *rpsH* | ECP_3394 | | | CTCATCTGTCT | -6 to +5 |
| JVpna-90 | *rplS* | ECP_2607 | | | CTCATAATTTA | -6 to +5 |
| JVpna-92 | *acpP* | ECP_1086 | | | CTCATACTCTT | -6 to +5 |
| JVpna-95 | scrambled | - | | | TTCACTATCTC | - |

**Dataset S1 (separate file).** This dataset includes (i) a list of all raw counts for each treatment condition for all biological replicates, (ii) a list of differential expression values for all identified transcripts in our RNA-Seq dataset involving at least three biological replicates, and (iii) the complete dataset for KEGG pathway analysis.
